# A sexually dimorphic hepatic cycle of periportal VLDL generation and subsequent pericentral VLDLR-mediated lipoprotein re-uptake

**DOI:** 10.1101/2023.10.07.561324

**Authors:** Tomaz Martini, Cedric Gobet, Andrea Salati, Jérôme Blanc, Aart Mookhoek, Michael Reinehr, Graham Knott, Jessica Sordet-Dessimoz, Felix Naef

## Abstract

Recent single-cell transcriptomes revealed spatiotemporal programmes of liver function on the sublobular scale. However, how sexual dimorphism affected this space-time logic remained poorly understood. We addressed this by performing scRNA-seq in the mouse liver, which revealed that sex, space and time together markedly influence xenobiotic detoxification and lipoprotein metabolism. The very low density lipoprotein receptor (VLDLR) exhibits a pericentral expression pattern, with significantly higher mRNA and protein levels in female mice. Conversely, VLDL assembly is periportally biased, suggesting a sexually dimorphic hepatic cycle of periportal formation and pericentral uptake of VLDL. In humans, *VLDLR* expression is also pericentral, with higher mRNA and protein levels in premenopausal women compared to similarly aged men. Individuals with low hepatic *VLDLR* expression show a high prevalence of atherosis in the coronary artery already at an early age and an increased incidence of heart attack.

## 1. INTRODUCTION

The mammalian liver is a vital metabolic organ, which processes both xenobiotics as well as nutrients, and supplies the body with carbohydrates, proteins and lipids based on recurring daily demands. The liver is the largest gland, and is composed of repetitive units called lobules, with hepatocyte layers organized around a central vein where deoxygenated blood exits. On the opposite side of the lobule, the bile duct, hepatic artery and portal vein form the portal triad. The artery and portal vein deliver oxygen and nutrients, respectively, resulting in nutrient and oxygen gradients along the periportal (PP) to pericentral (PC) axis, leading to differences in hepatocytes along the axis ^1^. In addition, the liver also shows striking daily physiological rhythms, which allow it to anticipate and respond to feeding-fasting and activity-rest cycles ^2^. Temporal programs are driven by recurring stimuli, such as nutrient availability, and by a genetically encoded molecular clock that drives self-sustained oscillatory processes called circadian rhythms. Taken together, the liver has a strong spatiotemporal organization of physiology ^3^, and perturbing this organization results in morbidity and mortality ^1,4,5^. However, it was not known how spatiotemporal functions were affected by sex beyond bulk and pseudobulk approaches ^6–9^. Sex strongly affects the incidence of cancers, including hepatocellular carcinoma (HCC) ^10,11^, susceptibility to drug adverse effects, including drug-induced liver injury (DILI) ^12^, as well as major cardiovascular events, with premenopausal women robustly protected against the latter ^10^. Moreover, sex differences in cardiovascular disease are amplified with increased dietary fat consumption ^10^. Very low density lipoproteins (VLDL), which supply tissues with triglycerides (TAG), and their remnants are the main culprits in atherosclerosis, and they are more atherogenic than low density lipoproteins (LDL) ^13–17^. VLDLs are produced by the liver and mature into VLDL remnants and LDL ^18^. In parallel, the liver can turn over a high amount of metabolites due to its size, and has a dominating role in lipid catabolism, enabling it to balance lipoprotein production and lipid degradation. Often, such opposing tasks in the liver are spatiotemporally compartmentalized ^1^.

Here we found that the hepatic VLDL receptor (VLDLR), critically involved in cardiovascular disease ^13,19–21^, is limited to the pericentral zone, and more highly expressed in females, with changes in its subcellular localization depending on the time of day. In parallel, a subset of genes involved in VLDL assembly show higher levels periportally. Electron microscopy (EM) enabled us to confirm that VLDL production shows a periportal bias (the process is more pronounced around portal veins), revealing a sexually dimorphic hepatic cycle of periportal formation and pericentral uptake of VLDL. In humans, *VLDLR* mRNA and protein levels were also increased centrally, and significantly higher in premenopausal women compared to men. Finally, individuals with low hepatic *VLDLR* levels showed a higher prevalence of atherosis in the coronary artery already at an early age and an increased incidence of heart attack.

## 2. RESULTS

### 2.1 Single-cell RNA sequencing reveals the space-, time- and sex-dependent organization of gene expression in the mouse liver

To obtain a first view into the aforementioned sexually dimorphic pathologies, intrinsically tied to the liver, we focused on sets of known pathology-associated genes (S. Table 1) and assessed their sex- and time-dependent expression from liver bulk RNA-seq (SFig. 1). Among the diseases considered, we found differences between the two sexes predominantly in genes associated with cardiovascular disease. Notably, there was higher female expression of *Vldlr*, while the low density lipoprotein receptor (*Ldlr*) and apolipoproteins (*Apo*s) showed comparable expression in both sexes (SFig. 1).

To investigate liver gene expression in function of time of day, lobular zone and sex, we performed droplet-based scRNA-seq of liver cells, collected, enriched for hepatocytes and cryopreserved at two time points, from male and female mice. The chosen time points, ZT10 and ZT22 (*Zeitgeber time*; ZT0 denotes when the lights are switched on, at ZT12 lights are turned off), correspond to physiologically distinct fasted (ZT10) and fed states (ZT22). Clustering of the cells revealed several nonparenchymal cell types as well as a main cluster of hepatocytes with an expected antizonated *Cyp2e1* (pericentral) and *Cdh1* (periportal) expression (Fig. 1A, SFig. 2A). Based on the clustering, we first assessed effects of time and sex on individual cell types in the pseudobulk, showing both global and cell-type specific regulation (SFig. 2B).

**Figure 1.**
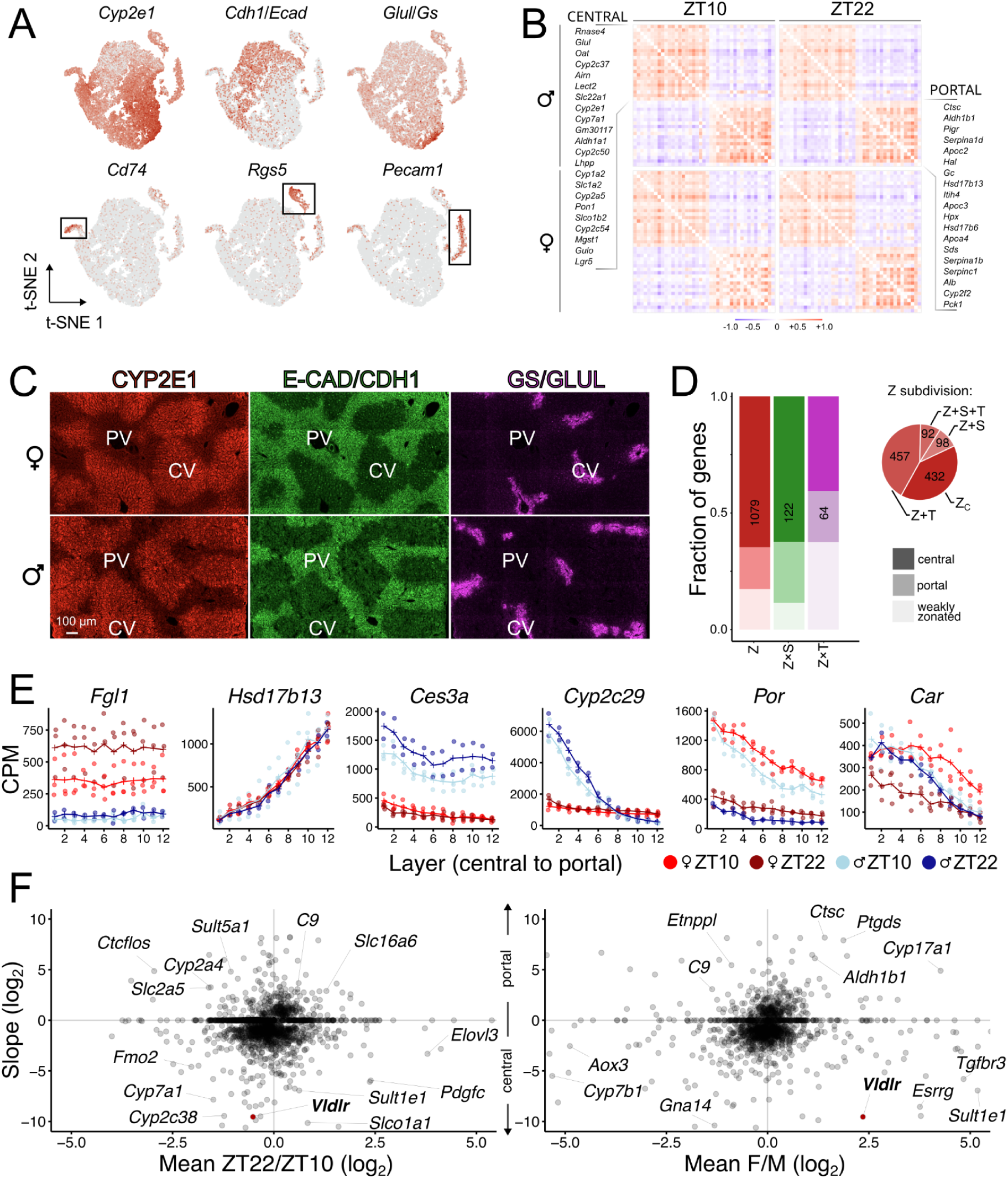
scRNA-seq uncovers cell types and the space-, time- and sex-dependent organization of gene expression in the mouse liver. (A) t-SNE plots show a main cluster of hepatocytes with antizonated Cyp2e1 and Cdh1/Ecad expression, and clusters of non-parenchymal cells identified with marker genes including Rgs5 (stellate cells), Pecam1 (endothelial cells), and Cd74 (immune cells; n = 12). (B) A Pearson’s correlation analysis reveals that the correlation between the selected portally and centrally biased genes is conserved independently of time and sex in hepatocytes (n = 12). (C) Immunofluorescence demonstrates that spatial patterns of canonical zonation markers E-CAD (periportal), and CYP2E1 and GS/GLUL (both pericentral) are indistinguishable in male and female mice. (D) Spatial reconstructions of lobular mRNA profiles combined with a model selection identified genes differentially affected by time (T), sex (S) and space (Z). The × denotes interacting effects (non-additive in log), resulting in changes of the shape of the spatial profile in function of either time or sex. The group of zonated genes (Z) also includes genes that change their mean expression levels with sex or time, but not the shape of zonation (in log), and can be subdivided (pie chart) into genes whose expression is > 50% higher in one sex (Z + S), at one time (Z + T), in function of both sex and time (Z + S + T), or constant across conditions (Z_c_). Most zonated genes have a pericentral bias. (E) Reconstructed spatial profiles of gene expression in 12 layers, 1 corresponding to the most pericentral layer. Counts are normalized to the total counts of cytoplasmic unique molecular identifiers (UMI) and represented as counts per million (CPM; n = 11; n = 2-3, per condition). Examples include a non-zonated gene (Fgl1; fibrinogen-like 1) and zonated genes, with the slope either not affected by time or sex (Hsd17b13), or with different effects of time and/or sex on the spatial profile in Ces3a (carboxylesterase 3a), Cyp2c29, Por (cytochrome P450 oxidoreductase) and Car/Nr1i3 (constitutive androstane receptor). (F) Sexual dimorphism and temporal changes of zonated genes, represented by the average slope (negative slopes characterize central profiles) across all conditions vs. the ZT22 to ZT10 (left panel) or female to male expression ratio (right panel). Genes in the corners are highly zonated and strongly regulated by time or highly zonated and sexually dimorphic, including Vldlr. Representative genes of functions described in the main text, and genes from S. Table 1 are explicitly named.

We identified two populations of hepatocytes that differed in the fraction of mitochondrial reads. Hepatocytes with low mitochondrial reads (< 3%) differed markedly in their overall gene expression (SFig. 3) and were therefore not included into the spatial reconstruction. These cells exhibited a downregulation of secreted protein genes, and an upregulation of ribosomal proteins, cytochrome o oxidases, *Jun*, and genes involved in gluconeogenesis ^22,23^. Such *mt-Co1*-negative hepatocytes were suggested to represent the liver stem cell niche ^24^.

Using the canonical hepatocytes, we performed a spatial reconstruction based on a set of established ^3,25,26^ and newly found zonation markers, which exhibited conserved correlation patterns across conditions (Fig. 1B, SFig. 4), as expected for a robust set of markers. To further confirm that zonation is not globally changed in males and females, we showed with immunofluorescence that the canonical parenchymal zonation markers CYP2E1, E-CAD and GLUL exhibited equivalent spatial expression patterns between males and females (Fig. 2C). To position each cell along the central to portal axis, we developed a maximum likelihood approach to infer 1D lobular coordinates directly from the scRNA-seq counts (S. Table 2). This revealed a variety of zonated expression profiles, including ones showing highly robust patterns across conditions, while others exhibited strong differential time and sex effects (Fig. 1D-E, S. Table 3). We categorized genes into nonzonated or zonated, with the latter accounting for approximately a fifth of the detected genes. These were additionally subcategorized based on effects of sex and time on the spatial expression profiles. More zonated genes showed a pericentral bias (Fig. 1D; SFig. 4-5). While sex and time often influence the mean expression levels of zonated genes ^7^, including metabolic and pathology associated genes (SFig. 5-6), we found that sex or time can also affect the shapes of zonation profiles (Z × S and Z × T genes, Fig. 1D).

**Figure 2.**
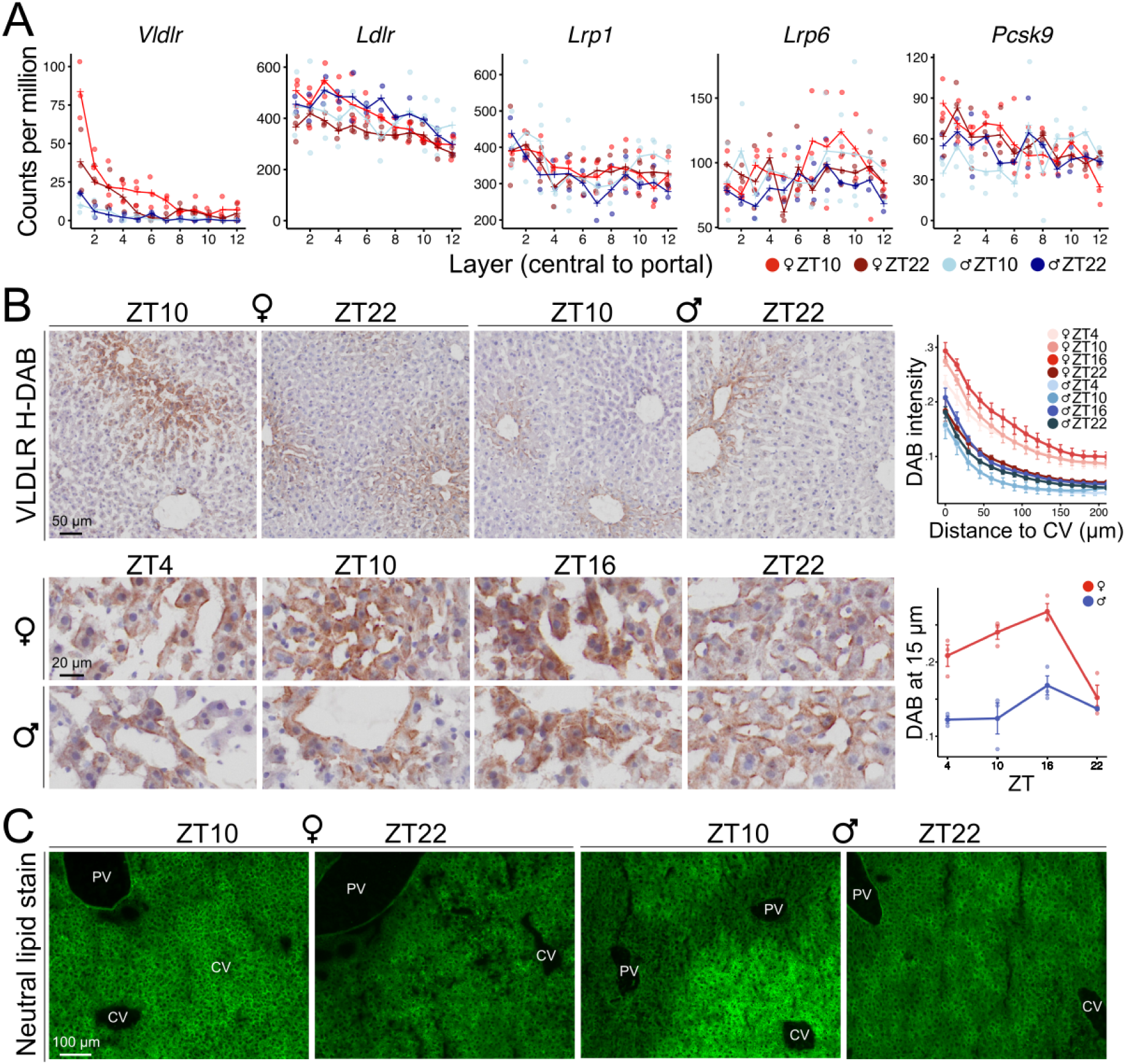
Pericentral and sex-dimorphic Vldlr expression in the mouse liver. (A) scRNA-seq revealed a sexually dimorphic and pericentral expression of Vldlr. In females, the expression is lower at ZT22. Other lipoprotein receptors, Ldlr, Lrp1 and Lrp6, do not exhibit sexual dimorphism (n = 11; n = 2-3, per condition). (B) H-DAB stainings on flash frozen tissue show a pericentral confinement of VLDLR, with more intensive immunoreactivity in females. Male and female immunoreactivity is cytoplasmic at ZT10 and ZT16 while more membrane-localized at ZT22 and ZT4. Quantification of DAB staining shown as relative signal in concentric areas of 15 µm around the central vein (CV), or quantification of DAB signal at the distance of 15 µm from the central vein at different time points, errors represent SEM between biological replicates (n = 24; n = 3, per condition). (C) Staining of neutral lipids with BODIPY 493/503 shows a pericentral increase in neutral lipids in males at ZT10, fitting with the space-time logic of cytoplasmic VLDLR immunoreactivity. Such a phenotype is not observable in females. At ZT22 there is a lack of zonation in the BODIPY signal (n = 2-3, per condition).

Focusing on sexual dimorphism in zonated metabolic and disease related genes, we identified many zonated genes showing highly sex-dimorphic mean expression levels (Fig. 1F, SFig. 5). Specifically, we observed major effects of sex on cytochromes (*Cyp*s, SFig. 7), solute carriers (*Slc*s, SFig. 8) and sulfotransferases (*Sult*s, SFig. 9). The strongly sex-^6^ and time-dependent ^27^ zonation profiles of *Cyp*s point to spatiotemporally compartmentalized programmes of xenobiotic biotransformation and drug-induced toxicity ^1^, which differ in male and female mice. Additionally, *Cyp7a1*, crucial in the neutral pathway of bile acid synthesis, varies with time, and *Cyp7b1*, essential in the acidic pathway of bile acid synthesis, is more highly expressed in males (SFig. 7). Solute carriers, some of which are associated with the sexually disparate DILI ^12^, show sex-dimorphic and time-of-day regulation, including the male-enriched and pericentral *Slco1a1* (SFig. 8). Intriguingly, we found with scRNA-seq that the aforementioned *Vldlr* was not only more highly expressed in females, but also pericentrally localized in both sexes (Fig. 1E; SFig. 5).

### 2.2 Sexually dimorphic, compartmentalized Vldlr expression and VLDL generation

VLDL and remnants are atherogenic ^13–17^, and VLDLR’s role in their trafficking is well established ^19–21,28,29^. In particular, hepatic VLDLR regulates circulatory TAG levels, as demonstrated in mice that expressed *Vldlr* exclusively in the liver ^28^. Furthermore, the upregulation of hepatic *Vldlr* is atheroprotective in mouse models of atherosclerosis ^19,20,30,31^. However, this upregulation was often argued to represent ectopic expression, because liver *Vldlr* levels were believed to be low ^19^, based on bulk tissue studies predominantly performed on male subjects. Hence, not much further attention was placed on hepatic *Vldlr*, despite significant hepatic VLDL uptake in rodents ^32–35^ and the observation that VLDL-sized lipoproteins were effectively cleared from the circulation even in *Ldlr* KO mice ^36–38^.

Strikingly, we found that *Vldlr* mRNA levels, previously reported to be very low in bulk liver ^28^, were markedly higher in pericentral hepatocytes and in female mice (Fig. 2A). Conversely, *Ldlr* expression was comparable in both sexes and not restricted to a particular sublobular zone. Immunostained sections of fresh frozen livers, isolated every 4 h, confirmed that VLDLR is pericentrally confined, with much stronger immunoreactivity in female tissue (Fig. 2B; SFig. 10-11). We also observed a pronounced daily rhythm in the subcellular localization of VLDLR in both sexes. Namely, cytoplasmic immunoreactivity was observed at the fasting-feeding transition, while membrane localization of the VLDLR signal was observed near the feeding-fasting transition (Fig. 2B). The results suggest that the uptake of lipoproteins by VLDLR is limited in time and space. We found that this patterning can be perturbed with the commonly prescribed fenofibrate (SFig. 10-11), a drug which functions through VLDLR-mediated hepatic TAG uptake ^28^. The perturbed spatiotemporal compartmentalization ^1^ of lipid metabolism could explain why fenofibrate leads to hepatic biochemical abnormalities in 5%-10% patients and complications such as DILI ^39^. Moreover, fenofibrate’s effects on VLDLR expression differed between females and males (SFig. 10-11), with the relative increase in VLDLR upon treatment generally larger in males, possibly explaining sex-dependent outcomes of fenofibrate treatment ^40,41^.

On the organismal level, intravenously injected DiI-labeled VLDL particles accumulated in the liver (SFig. 12), fitting with previous reports of significant hepatic VLDL uptake in rodents ^32–35^, further supporting a central role of the liver in lipoprotein catabolism^19^. Since females have a higher hepatic *Vldlr* expression, we hypothesized that this may increase the clearance of VLDL and remnants from the blood of female mice. A higher clearance model would suggest lower concentrations of circulatory TAGs in females, which is indeed observed in bulk lipidomics ^42^ data (SFig. 13). On the sublobular level, we saw an increase in neutral lipids (TAGs and cholesteryl esters) around central veins at ZT10 in males (Fig. 2C, SFig. 14), consistent with the pericentral expression of VLDLR and the temporal dynamics of cytoplasmic VLDLR immunoreactivity, indicative of lipoprotein internalization. This pericentral bias in neutral lipids was attenuated in females, possibly due to lower levels of circulatory lipoproteins and/or increased lipid catabolism (e.g. *Lipa*, *Lipc*, *Mgll*) following lipoprotein internalization (SFig. 5, S. Table 2).

Contrary to pericentral *Vldlr*, *Apoe* and *Apoa4*, the latter involved in TAG secretion and VLDL expansion ^43^, as well as *Tm6sf2*, involved in TAG secretion via lipidation of VLDL ^44–46^, show higher periportal levels and higher expression in females in the scRNA-seq. *Apoc2* and *Apoc3*, found on VLDL particles, were also higher periportally. Both APOA4 and APOC2 are physiologically important for recognition by the lipoprotein lipase (LPL) and its enzymatic activity, allowing LPL to hydrolyze TAGs carried by lipoproteins ^47–49^. Thus, we identified a periportal bias of genes involved in VLDL generation, though *Apob*, lipidated to form VLDL, does not follow that pattern, but shows attenuated expression in the midlobular region (Fig. 2A). Of note, *Hsd17b13*, described as a protein associated with lipid droplets ^50^, shows robust periportal expression (Fig. 1E), and the leptin receptor (*Lepr*), which affects generation of VLDL particles ^51^, also shows a periportal bias in females, with much higher female and ZT10 expression (SFig. 6). Given the possibility that pre-VLDL fuses with lipid droplets to form VLDL-sized particles ^52^, *Hsd17b13*’s periportal expression is intriguing.

To assess the implications of these zonated transcript levels on the spatial properties of VLDL assembly, we employed transmission electron microscopy (TEM) on livers from 3 female and 3 male mice at ZT2. This time was antiphasic to the cytoplasmic VLDLR immunoreactivity and coincides with the predicted postprandial VLDL assembly ^53,54^. Our optimized TEM stainings allowed us to visualize VLDL-containing vesicles, localized at the Golgi, which were abundant periportally, but scarce pericentrally in both sexes (Fig. 3B, SFig. 15). The identification of these VLDL-containing vesicles was based on their electron density in TEM and well-established specific morphology ^53,55^, as well as on their known pathway of assembly and subcellular localization ^56,57^. For quantification of VLDL-containing vesicles in 3D cellular volumes, we employed scanning electron microscopy (SEM), showing a 2-fold periportal to pericentral ratio of vesicles loaded with VLDL particles (Fig. 3C). These data indicate a periportal generation of carriers loaded with VLDL particles, consistent with spatial patterns of gene expression, demonstrating spatial compartmentalization of VLDLR-mediated lipoprotein catabolism and VLDL generation. EM also revealed differently shaped periportal mitochondria, which we reconstructed in 3D, fitting with a zonated mitochondrial transcript fraction observed previously ^3^ and found here (SFig. 16).

**Figure 3.**
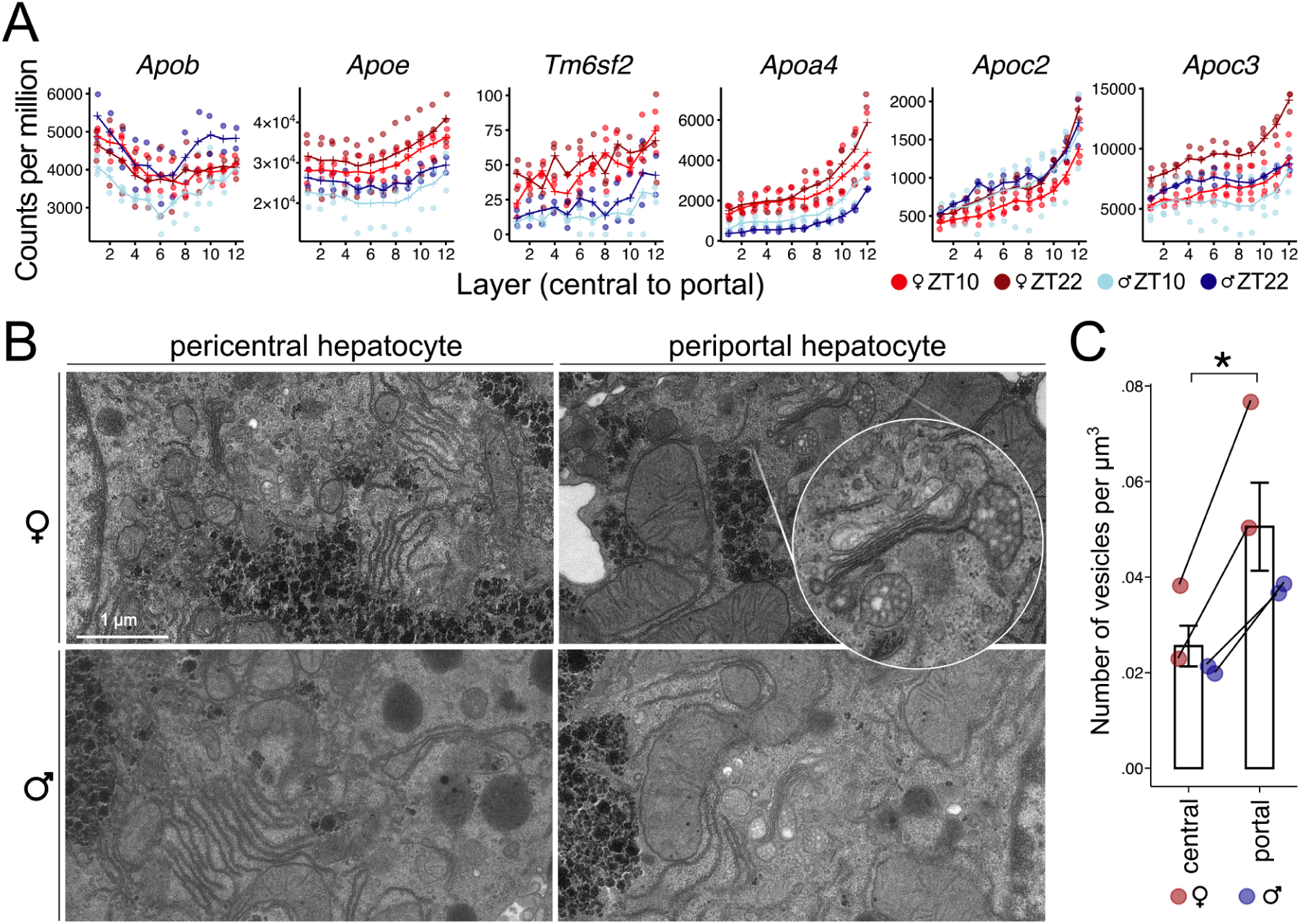
Periportal VLDL generation in the mouse liver. (A) scRNA-seq revealed a periportal bias of multiple genes crucial for VLDL generation. Apoe, Apoa4, Apoc2, Apoc3 and Tm6sf2 show a periportal bias. Apob shows attenuated mRNA counts in the midlobular region (n = 11; n = 2-3, per condition). (B) Transmission EM images of periportal and pericentral hepatocytes. Periportal hepatocytes are rich in vesicles loaded with low density particles, whereas pericentrally these vesicles are scarce in both females and males. The scale bar represents 1 µm (n = 6 mice; n = 3, per sex). (C) Scanning EM allowed quantification of vesicles loaded with VLDLs in > 40000 µm^3^ of pericentral and periportal hepatic tissue, revealing an approximately 2-fold periportal to pericentral ratio of VLDL-loaded carriers (n = 4 mice; p = 0.017, paired t-test).

### 2.3 VLDLR levels in humans recapitulate pericentral confinement and higher abundance in women

To investigate whether our findings are conserved in humans and to address the controversial low hepatic *VLDLR* ^19,28^, we assessed *VLDLR* mRNA levels in 40 tissues of human donors from the Genotype-Tissue Expression (GTEx) consortium ^27,58^. Additionally, we compared VLDLR protein abundance across 21 tissues ^59^. While the bulk liver showed approximately 4-fold lower *VLDLR* mRNA compared to most assessed tissues, on the protein level, hepatic VLDLR levels were comparable to those in other organs, with the exception of the heart, where its abundance was approximately 5-fold higher (Fig. 4A, SFig. 17). Protein levels of VLDLR in the bulk were comparable to those of LDLR, which has a well-established physiological role in the liver. Importantly, VLDLR’s spatial confinement would imply even higher mRNA and protein levels in central hepatocytes.

**Figure 4.**
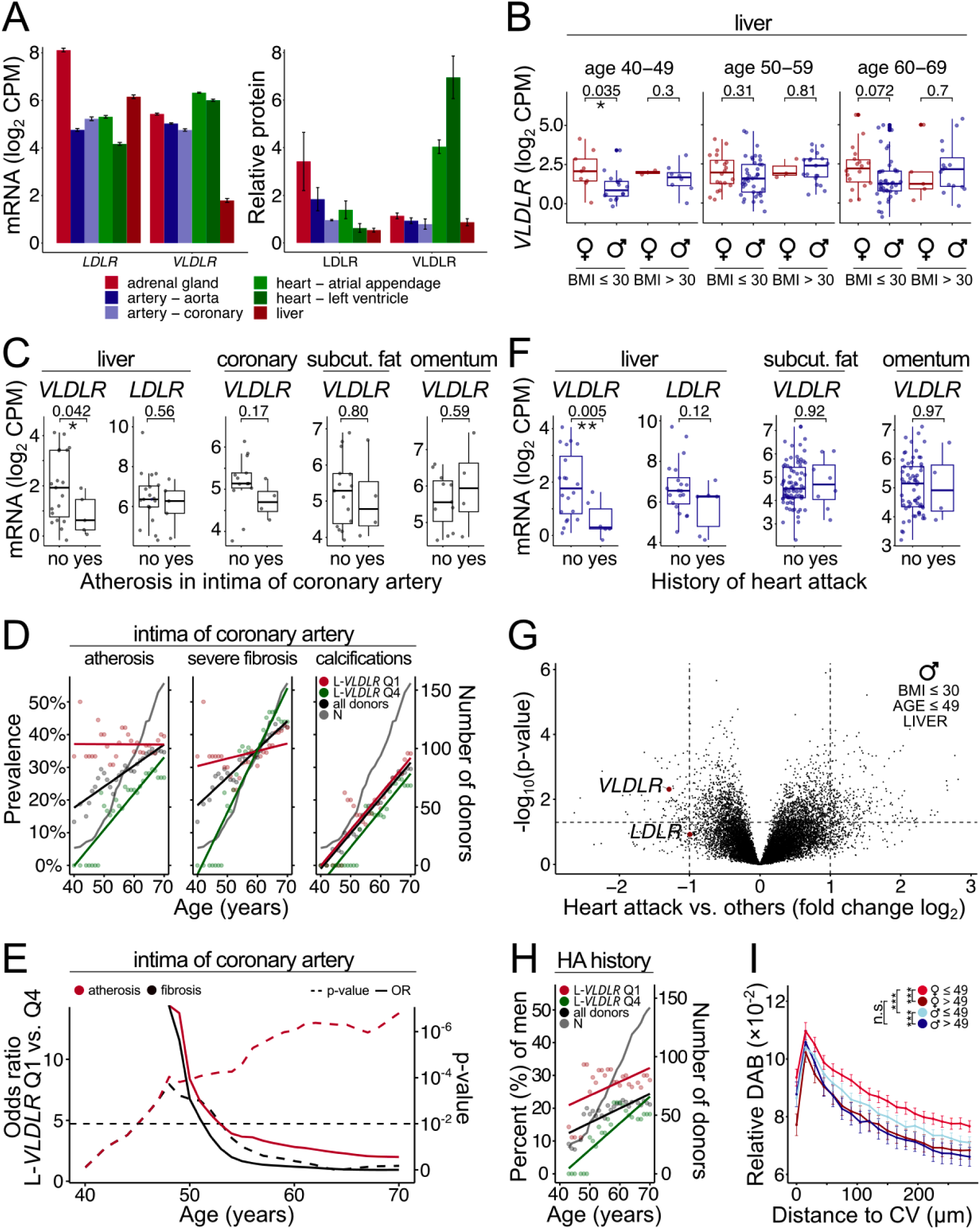
VLDLR expression in human tissues. (A) Mean mRNA levels from bulk RNA-seq (left panel; error bars represent SEM) and relative mean protein levels from bulk proteomics (right panel; reanalysis of data from ref. 59; error bars represent SEM) in human tissue. While hepatic VLDLR mRNA levels are lower, the VLDLR protein abundance is comparable to that of LDLR in the liver. Hepatic VLDLR protein abundance is comparable to that in the aorta and coronary artery. (B) Stratification of liver bulk RNA-seq reveals 88% higher VLDLR mRNA levels in 40-49 year-old premenopausal women compared to age-matched men in the BMI ≤ 30 group, or 122% higher after outlier removal (more than 3 standard deviations from the group mean). In obese individuals with a BMI > 30, this difference is lost. The center line represents the median, error bars represent the interquartile range, p-values were calculated with a t-test, without outlier removal. (C) In young (≤ 49 years) non-obese (BMI ≤ 30) men and women, intimal atherosis correlates with lower hepatic VLDLR mRNA, but not with hepatic LDLR or adipose tissue VLDLR. (D) All donors, regardless of age or BMI, were categorized based on the quartile of liver VLDLR mRNA. The fraction of individuals below a certain age with a specific condition was computed for the lowest quartile (Q1; red line), the highest quartile (Q4; green line), and the overall donor group (black line). For easier visualization, a linear fit across the data-points is shown. (E) The comparison of Q1 to Q4 donors shows that the odds ratio for developing intimal atherosis is significantly higher in the Q1 group across all ages or until about age 50 for fibrosis. At younger ages the odds ratio is large due to the absence of donors within Q4 with these manifestations. (F) Young (≤ 49 years) non-obese (BMI ≤ 30) men were further stratified based on an annotated medical history of heart attack. Those without such a medical history show higher hepatic VLDLR mRNA levels, but there were no differences in adipose tissue VLDLR or hepatic LDLR between the two groups. (G) Assessment of genome-wide differences in hepatic gene expression in young non-obese males with vs. without a history of heart attack identifies VLDLR among the most differentially regulated and statistically significant hits. (H) All male donors, regardless of age or BMI, were categorized based on the quartile of liver VLDLR mRNA. A higher percent of men within the Q1 group experienced a heart attack compared to those in the Q4 group. (I) Quantification of immunohistochemistry shown as mean relative DAB intensity per group in function of distance from the central vein (CV). Donors were grouped by age and sex as shown in the legend (n = 31; error bars represent SEM). The staining in the groups of men and women > 49 years are potentially overestimated due to the presence of the age pigment lipofuscin.

Next, we stratified the expression of *VLDLR* by sex, age and body mass index (BMI). Hepatic *VLDLR* mRNA was higher in premenopausal (40-49 years) female donors compared to males (Fig. 4B). The difference was lost in obese and in the postmenopausal (> 49 years) and age-matched individuals. There were no differences between the groups in *APOB* or *APOE*, but there was a trend of higher *LDLR* levels in young men (SFig. 18), possibly due to their attenuated ability to internalize larger VLDL-sized particles ^36^, prolonging their half-life and promoting their maturation into LDL.

Given VLDLR’s established central role in lipoprotein internalization ^19,20,60^ and the abundant hepatic clearance of atherogenic lipoproteins in rodent studies ^19^, we assessed whether hepatic *VLDLR* levels correlate with a risk for atherosclerosis or myocardial infarction in humans. Based on our clinical experience, the coronary artery is an ideal tissue for estimation of atherosclerosis, as atherosclerosis foci are often distributed along larger coronary artery segments and frequently also occupy larger contiguous areas. This in turn results in a lower sampling error. Hence, to assess potential links between hepatic *VLDLR* and atherosclerosis, a cardiovascular pathologist evaluated coronary artery H&E stainings from 179 donors who had corresponding GTEx hepatic bulk sequencing data (S. Table 4).

Since atherosclerosis is a progressive disease that typically develops over many years, studying young individuals can help identify factors that contribute to early-onset atherosclerosis and minimize confounding factors that affect overall health over time. Young individuals are usually also less medicated, whereas in older people the use of antilipemics is prevalent. Hence, we first assessed this age group. Young non-obese donors with atherosis in the intima had significantly lower hepatic *VLDLR* expression compared to those without intimal atherosis, but exhibited no differences in hepatic *LDLR* or adipose *VLDLR* (Fig. 4C). Next, we assessed the prevalence of pathological changes in the coronary arteries of all donors from the lowest (Q1) and highest quartile (Q4) of hepatic *VLDLR* expression in function of age (Fig. 4D). We found that the prevalence of intimal atherosis and severe fibrosis, both associated with arterial lipid accumulation that leads to inflammation and atherosclerotic plaque formation, was low in the Q4 group in young individuals and then significantly increased with age, while it was already high in young Q1 donors (Fig. 4D). However, we saw no association of hepatic *VLDLR* with calcifications of the coronary artery, which are manifestations that are not associated with lipoproteins. Strikingly, the prevalence of intimal atherosis in donors within the lowest quartile of liver *VLDLR* mRNA was already 40% at the age of 40 years. Contrary, for the same age, the prevalence was 0% for those in the highest quartile of hepatic *VLDLR*. Notably, Q1 and Q4 appear to intersect at approximately 70 years of age. Importantly, hepatic *VLDLR* mRNA levels affect the odds ratio for lipid deposition within the coronary artery (Fig. 4E).

Since advanced stages of atherosclerosis can lead to a heart attack, we evaluated whether hepatic *VLDLR* levels correlate with such events. In young (≤ 49 years) non-obese (BMI ≤ 30) men, lower hepatic *VLDLR* levels significantly correlated with an annotated medical history of heart attack, myocardial infarction or acute coronary syndrome (henceforth heart attack; Fig. 4F, SFig. 19). GTEx donors with such a history showed 2.5-fold lower mean (2.8-fold median) hepatic *VLDLR* mRNA levels. In comparison, there was no significant association between these instances and hepatic *LDLR* expression. Likewise, there was no correlation of heart attack history with adipose tissue *VLDLR* (Fig. 4F, SFig. 20). Genome-wide, in donors with a history of heart attack, *VLDLR* in the liver stands out among the most dysregulated genes (Fig. 4G), while other tissues do not show a similar association (SFig. 20). In females, the sample size for a medical history of heart attack was too low for assessment, as premenopausal women are robustly protected against coronary heart disease due to lower circulating TAG and APOB levels ^61,62^ and a higher clearance of circulating VLDL-associated TAGs ^18,63^. To overcome the constraint of the small sample size of young male postmortem tissue donors with a history of heart attack, we also assessed how the percent of men that experienced a heart attack changes with age for donors within the lowest (Q1) and highest (Q4) quartile of liver *VLDLR* mRNA expression, using the full dataset. The percent of men with a heart attack was substantially higher in those individuals with low hepatic *VLDLR* expression (Fig. 4H).

At the sublobular scale, immunostaining of formalin-fixed paraffin-embedded (FFPE) sections from 33 human donors (S. Table 5) confirmed a pericentral localization of VLDLR (Fig. 4I; Fig. 5; SFig. 21-24). Corresponding H&E stained slides were assessed by a clinical gastrointestinal pathologist to exclude the presence of significant macrovesicular and microvesicular steatosis. In a subset, a prominent pericentral deposition of lipofuscin was present. To exclude the possibility that a significant part of the signal in the VLDLR immunohistochemistry was due to lipofuscin, additional immunohistochemical stains using a purple chromogen were performed (SFig. 24). We observed a generally higher premenopausal women VLDLR immunoreactivity compared to age matched men (Fig. 4I, SFig. 23). Similarly to mRNA, the difference was lost between the postmenopausal and male age-matched donors, and intraindividual variability was more apparent in this age group. Highest protein VLDLR levels were observed in premenopausal women, and these dropped after menopause (SFig. 23), fitting with observed increases in circulating VLDL, LDL and TAGs in postmenopausal women ^64,65^.

**Figure 5.**
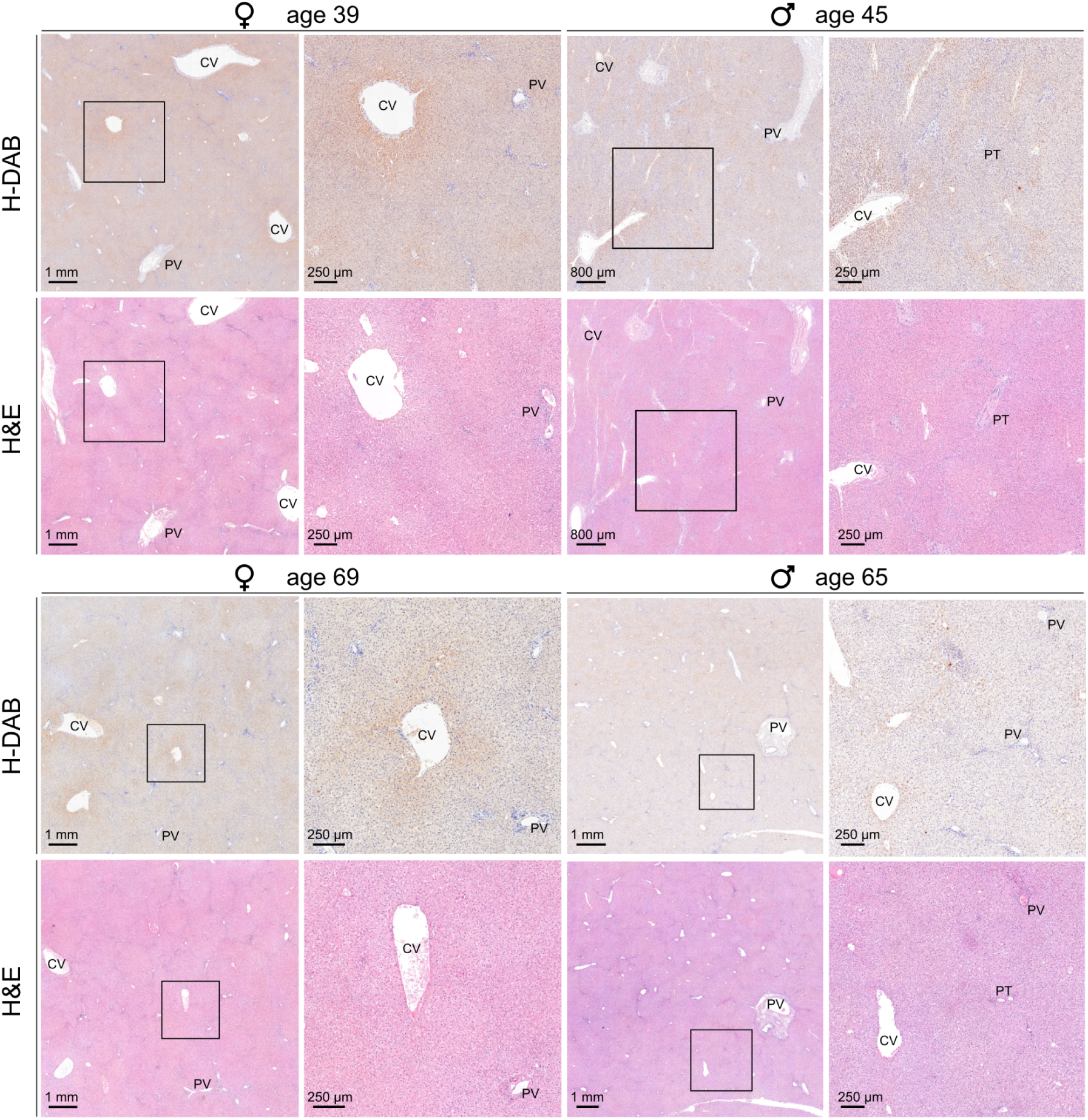
VLDLR immunoreactivity in human liver tissue. H-DAB VLDLR immunostainings and corresponding diagnostic H&E sections from the same FFPE block. Samples were grouped by sex and age, with the younger group consisting of premenopausal female donors and age matched males (≤ 49 years). Based on the quantified DAB signal, non-pathological samples with the median immunoreactivity were chosen for representative images. For each representative sample, a lower magnification overview and a higher magnification image are shown. Squares mark the magnified area. The H-DAB stainings show VLDLR immunoreactivity around the central veins in all samples. The highest DAB stain intensity is seen in premenopausal females (n = 33 donors; CV - central vein, PV - portal vein, PT - portal tract).

Taken together, the mouse experiments allowed us to identify a pericentral expression of hepatic VLDLR, which was conserved in humans. VLDLR mRNA and protein levels were higher in young women compared to men, and *VLDLR* mRNA levels correlated with atherosclerosis and heart attack incidence.

## 3. DISCUSSION

We uncovered higher hepatic VLDLR abundance in female mice and young women compared to males, with central sublobular domains of VLDLR immunoreactivity. Unlike earlier work suggesting that VLDL generation occurs pericentrally ^66^, our data indicate that VLDL generation likely occurs periportally after feeding ^54^ to sustain the organism during its fasting phase. Conversely, the subsequent pericentral hepatic re-uptake of circulating lipoproteins by VLDLR occurs primarily at the end of this fasting phase.

The blood levels of VLDL, remnants and LDL as well as APOB and TAG are lower in women than in men, and this is believed to be a major factor in the sexual dimorphism of coronary heart disease ^10,62,67^. Additionally, plasma clearance of VLDL-associated TAGs is much higher in women than men ^18,63^. However, the underlying cause of this sexual dimorphism remained elusive. Given the well established role of hepatic VLDLR in the internalization of lipoproteins and its atheroprotective role in rodent studies, it is reasonable to assume that the increased hepatic VLDLR expression in females could play a role in the marked differences in circulatory lipoprotein and TAG levels between young women and men, or even contribute towards the sexual dimorphism observed in cardiovascular disease. While further research into the association of hepatic VLDLR and cardiovascular disease is needed, our human studies indicate that low hepatic *VLDLR* mRNA levels correlate with the incidence of heart attack in young men. Moreover, low hepatic *VLDLR* expression strongly correlates with coronary artery intimal atherosis, whose prominent manifestation is the buildup of fats in the artery wall. Thus, evidence converges to suggest that hepatic *VLDLR* is critically involved in lipoprotein homeostasis. Hence, it will be important to assess the role of pericentral VLDLR, as well as potential sex differences in downstream lipid catabolism, in a wider (patho-)physiological context, especially in metabolic dysfunction-associated liver disease (MAFLD), which is marked by a pericentral hepatic lipid accumulation. Importantly, the liver is ideally positioned for lipoprotein catabolism, as its large size enables a considerable metabolite turnover, while its discontinued capillaries allow large lipoproteins to come into contact with the hepatic parenchyma.

Pharmacologically, systemic TAG levels are lowered by fibrate administration, which are commonly prescribed PPARɑ agonists. Fenofibrate in mice reduces circulating TAGs through the upregulation of hepatic *Vldlr*, thereby increasing hepatic lipoprotein intake. This effect is achieved by activating *Vldlr*’s transcriptional regulator PPARɑ ^21,68–70^. Fibrates have shown success in some, but fallen short of expectations in other clinical trials ^71^, with different outcomes between men and women ^41^. This calls for a more personalized approach, as some patients might benefit from fibrate-mediated *VLDLR* upregulation and others from alternative strategies, including the attenuation of VLDLR degradation with the emerging class of PCSK9 inhibitors ^72^. Speculatively, *VLDLR* expression could already be closer to the plateau in certain groups, resulting in limited success of fibrate administration. Importantly, fenofibrate and PCSK9 inhibitors act on the hepatic re-uptake phase of the VLDL cycle, while statins, lomitapide, ezetimibe and bile acid sequestrants act on lipid availability for VLDL generation, which should be taken into consideration when designing therapeutic strategies.

Whereas previous research already indicated a possible significance of hepatic VLDLR in the systemic regulation of the most atherogenic lipoproteins, those results were viewed with reservation due to a general belief that hepatic *VLDLR* expression was low or even absent ^19^. Hence, our discovery that the pericentrally expressed *VLDLR* mRNA is diluted in bulk tissue and highly sexually dimorphic sheds light on a previously overlooked aspect of VLDLR biology and proposes an explanation for the sexual dimorphism in lipoprotein homeostasis, based on a hepatic cycle of periportal formation and sex-dimorphic pericentral lipoprotein re-uptake.

## Acknowledgements

We thank Stéphanie Clerc-Rosset for TEM imaging, Olivier Burri for producing a script for DAB immunostaining quantification, Gian-Filippo Mancini for histological processing of tissues, Roy Combe for whole organ fluorescence imaging, Irene Centeno-Ramos for coordinating experiments on human tissue, and Christine Göpfert and Giovanni D’Angelo for advice in their respective fields of animal pathology and lipidomics. The work was supported by the Swiss National Science Foundation project grant 310030B_201267 to F. N.

## Author contributions

T. M. and F. N. conceived and designed the experiments. T. M. performed most of the experiments, analyzed the data and prepared the figures. C. G. and A. S. performed the computational analysis, A. M. assessed the human liver tissue, while M. R. assessed the coronary artery samples. A. M. and M. R. provided their clinical expertise. J. B. and G. K. planned and analyzed electron microscopy experiments. J. S. D. planned and oversaw the histology. T. M. and F. N. wrote the manuscript.

## Data availability

All scRNA-seq data have been deposited in GEO with accession code GSE244392.

## Code availability

The codes that support the findings of this paper are available from the corresponding author upon request.

## 4. MATERIALS AND METHODS

### Animal experimentation

All animal experimentation was done according to local laws, and approved by the veterinary office of the Canton of Vaud, Switzerland. Mice were housed under a 12 h light-dark cycle and temperature of 21 ± 2 °C, with ad libitum access to food and water. The time of a perturbation is denoted with *zeitgeber time* (ZT), in accordance with the nomenclature in chronobiology, and ZT0 and ZT12 refers to the time of the switching on or off of lights, respectively. Experiments were performed with 129S6/SvEv (Taconic) mice, abbreviated as 129S in the manuscript. Additional and comparative data from C57BL/6 and CD-1 strains came from previous work, as described in corresponding method and result sections.

### Handling of human tissue

All human tissue handling, storage and human experimentation was performed according to local laws, and the work was approved by the Cantonal commission of Vaud for ethics in human research. All participants signed a written consent form. The tissue (S. Table 5), assessed by a clinical gastrointestinal pathologist (A. M.), was obtained from the Tissue Biobank Bern (TBB) of the University of Bern, with the criteria to exclude steatotic tissues. The TBB sectioned FFPE at 4 µm and flash frozen samples at 10 µm thickness. Further manipulation of the tissue was performed in house, except H&E staining and scanning, which was done at the University of Bern.

### Analysis of mouse bulk sequencing data

C57BL/6 RNA-seq data was reanalysed from Weger et al. ^7^. Differential expression analysis was previously performed using DESeq2 ^73^.

### Analysis of human mRNA data (GTEx RNA-seq)

Raw gene read count data was obtained from GTEx V8 (2017-06-05_v8_RNASeQCv1.1.9, dbGaP Accession phs000424.v8.p2). For each tissue, genes with fewer than 10 counts across the samples were excluded. Counts were normalized based on library size and scaled using the edgeR TMM method ^74^, followed by conversion to counts per million (CPM). After the addition of a pseudo-count of 0.25, CPM values were log-transformed (log_2_). Information on sex, age, and BMI was retrieved from the dbGap metadata (phs000424.GTEx.v8.p2.c1.GRU) for every donor. The history of heart attack and acute coronary syndrome was taken from the metadata category MHHRTATT. Differential expression was assessed between age, sex and BMI groups using a t-test.

### scRNA-seq

#### Isolation and cryopreservation of cells

For perfusions, a peristaltic pump was used to withdraw buffered solutions from a water bath, kept at 42 °C, via a system of medical grade silicone tubing. The tubing led to a Graham condenser, which was heated with a circulating water flow from the water bath, utilizing a separate pump. The solutions exited the Graham condenser vial medical grade silicone tubing, which was connected via Luer adapters to a butterfly needle. The system ensured that all solutions had an approximate temperature of 37 °C at the tip of the butterfly needle. To perform the perfusion, mice were euthanized with 150 mg per kg pentobarbital. Upon absence of reflexes, the cadavers were opened and the butterfly needle, connected to the peristaltic pump system, was inserted into the inferior vena cava and clamped into place with a surgical hemostat. The perfusion pump was started and the portal vein was immediately cut to prevent an increase of pressure within the liver. First, to flush the blood, the perfusion was performed for 3 min using a buffer without collagenase. Subsequently, the mice were perfused for 3 min with a solution containing collagenase. Intact livers were isolated, placed into approx. 15-20 ml DMEM : F12, and cells released by breaking the Glisson’s capsule. Next, straining through a 70 µm mesh and enrichment for hepatocytes based on centrifugation (50 G for 5 min, with a low breaking speed) was performed. For the centrifugation, each sample was split in two: one falcon was immediately stored as described below. The other falcon was used for resuspension of the pellet, and cell counting and viability assessment (trypan blue), followed by another round of centrifugation. After isolation, we implemented a isopentane cell freezing protocol to preserve cells isolated at the two precise time-points, allowing us to prepare sequencing libraries at a later point. Briefly, after centrifugation, the whole falcon tube containing the pellet enriched for hepatocytes was dipped into isopentane, precooled to -80 °C. The cell viability assessment upon thawing (see below) confirmed the feasibility of the freezing protocol.

#### Preparation of libraries for single-cell sequencing

The cryopreservation reduced hands-on time significantly. It also enabled us to thaw the cells immediately before loading onto the microfluidics chip of the scRNA-seq kit, reducing cell sticking and appearance of doublets in the dataset. Briefly, cells stored in 50 ml falcon tubes were placed on ice, and DMEM : F12 (1 : 1) medium with 10% FBS, cooled at 4 °C, was immediately added to a final concentration of 2 × 10^6^ cells per ml. Within 5 min, the cells were gently resuspended with a 50 ml serological pipette. The addition of 10% FBS to the medium reduced cell aggregation and the larger serological pipette reduced shear stress on the cells. The cell concentration and viability was assessed and the cells were immediately mixed with the 10x Genomics reagents and loaded on a pre-prepared (glycerol already in empty rows) 10x Genomics Chromium chip. Library preparation was performed using the 10x Genomics Chromium v3 kit according to manufacturer’s instructions. The assessment of viability post freezing showed virtually no loss of cells.

### Analysis of mouse scRNA-seq data

#### Filtering and clustering

Raw sequencing data were aligned to the mouse genome (mm10) using CellRanger (6.1.2). Intron and exon counts were computed using Velocyto ^75^. From all sequenced GEMs (gel beads in emulsion), cells were selected on a per-sample basis by evaluation of the intronic fraction of individual GEMs (refer to ^76^). GEMs with no or low introns were removed for each sample. A cutoff between a low and mid intron distribution was set on a per-sample basis, but at 8.5% for most samples. As hepatocytes are highly metabolically active with a high percent of mitochondrial genes in the transcriptome ^77^, and nonparenchymal cells have lower mitochondrial fractions, a filter of < 40% mitochondrial reads was set. Initial analysis was performed using the Seurat 4.3.0 package. Briefly, samples were normalized (sctransform) and integrated (features = 3000) and the cells clustered based on a t-distributed stochastic neighbor embedding (t-SNE) dimensionality reduction and Louvain algorithm (resolution = 0.5). This allowed us to identify individual cell types. For quantifications of gene expression in identified clusters of individual celly types, non-integrated transcripts (UMIs) were normalized to the total cytoplasmic gene count (excluding mitochondrial genes) since the mitochondrial fraction changes in function of cell type and lobular zone in hepatocytes. We identified a subcluster of hepatocytes that separated from others by low expression of secreted proteins and which exhibited a low mitochondrial fraction. To split the hepatocytes into those with low and high mitochondrial fractions, a cutoff of 3% mitochondrial genes was chosen. For the high mitochondrial gene subpopulation of hepatocytes (≥ 3%), a lobular position was calculated (spatial reconstruction), as described below. The pseudo-bulk calculations were done on all samples, with 3 biological replicates per condition. For spatial reconstruction and subsequent plotting of expression profiles, one male ZT22 sample was removed due to overall lower UMI counts.

#### Spatial reconstruction of hepatocyte zonation profiles from scRNA-seq

We developed a probabilistic model to assign a one-dimensional spatial position to every cell based on the expression patterns of established zonation marker genes in individual cells ^3,25,26^. The initial set of markers, taken from ref. 22, included *Pck1*, *Aldh1b1*, *Ctsc*, *Sds*, *Hal*, *Hsd17b13*, *Cyp2f2* as portal and *Oat*, *Cyp2e1*, *Lect2*, *Cyp2c37*, *Gulo*, *Cyp2a5*, *Glul*, *Aldh1a1*, *Cyp1a2*, *Slc22a1*, *Slc1a2* as central markers. The spatial latent lobule coordinate *x* was determined by maximizing the likelihood of the observed raw counts, given *x* and other model parameters. We used a negative binomial noise model for the read counts to best capture the fluctuations of scRNA-seq data ^73^. The expectation of the negative binomial distribution for a cell *c* and gene *g* in sample *s* was modeled by µ*_gcs_* = *N_cs_exp*(*a_g_x_cs_* + *b_gs_*), where *a_g_* and *b_gs_* are gene-specific slope and intercept coefficients. Note that the gene slopes *a_g_* are the same across all samples, while intercepts *b_g,s_* can be sample dependent. Thus we assumed that there are landmark zonation genes with invariant slopes across all the samples, and these define the lobular coordinates. Finally, *N_cs_* is the number of unique molecular identifiers (UMI) captured for cell *c* in sample *s*. The dispersion parameter α was also fitted and it is unique across all samples and cells. The fitted values of the dispersion parameter (0.076 for the initial gene set and 0.096 for the enriched one) are quite low indicating minimal overdispersion, closely approximating a Poisson model.

This model displays inherent symmetries without further constraints on the parameter, in fact the model has rank-deficiency *s* + 1. One symmetry is a scaling invariance as the expected valuesg μ*_gcs_* only depends on the product *a_g_x_cs_*. We broke this symmetry by setting the slope *a_g_* of a selected gene (*Cyp2e1*) to 1 (which also sets the directionality of the central to portal axis, i.e. portal genes have a positive slope). The remaining *s* symmetries concern translational invariance: the shifting of all cells in a sample by the same quantity *q_s_* redefines the intercept *b’_gs_* = *b_gs_* + *a_g_q_s_*. To set the reference, we fixed *q_s_* such that cells expressing the highest central marker genes (e.g. *Glul*) were aligned across all the samples.

We initialized cell positions *x_cs_* using principal component analysis (PCA). Specifically, the projection on the first principal component (PC) was used. This choice was justified since it was clear that the first principal component correctly separated central and portally expressed genes. The initial gene parameters, *a_g_* and *b_gs_* were fitted to the PCA positions through negative binomial regression. Once we obtained these complete sets of initial values, we used gradient descent to optimize positions and gene parameters together.

In a second step, once the cell coordinates were estimated as described above, we expanded our set of reference zonation markers to enhance the robustness of the spatial assignment. Initially, we chose a subset of 400 genes that showed the most significant deviations from a linear trend in the coefficient of variation (CV) versus mean plot, as the initial set of marker genes exhibited substantial divergence from this trend. Next, we fitted the model’s slopes and intercepts for every gene, while keeping the cell positions *x* previously fitted with the smaller/initial set of genes fixed. Large positive or negative slopes, corresponding to central or portal genes, allowed us to identify potential zonation markers. Here to account for inter-mouse variability, we let the slope coefficient be sample dependent (μ*_gcs_* = *N_cs_exp*(*a_gs_x_cs_* + *b_gs_*)) which was not the case in the first model. Finally, we chose genes that had the highest slope’s mean to variance ratio (highest signal to noise) across the different mice. The final enriched set of genes included *Ctsc*, *Aldh1b1*, *Pigr*, *Serpina1d*, *Apoc2*, *Hal*, *Gc*, *Hsd17b13*, *Itih4*, *Apoc3*, *Hpx*, *Hsd17b6*, *Apoa4*, *Sds*, *Serpina1b*, *Serpinc1*, *Alb*, *Cyp2f2*, *Pck1* as portal and *Rnase4*, *Glul*, *Oat*, *Cyp2c37*, *Airn*, *Lect2*, *Slc22a1*, *Cyp2e1*, *Cyp7a1*, *Gm30117*, *Aldh1a1*, *Cyp2c50*, *Lhpp*, *Cyp1a2*, *Slc1a2*, *Cyp2a5*, *Pon1*, *Slco1b2*, *Cyp2c54*, *Mgst1*, *Gulo*, *Lgr5* as central markers.

The plotted expression profiles (Fig. 1-2) were generated by binning the 1D coordinates into 12 bins, representing layers of hepatocytes around the central vein. The expression is represented as counts per million (CPM), normalized by the total gene count excluding mitochondrial genes. The expression profiles can be generated using data from S. Table 2.

#### Model selection on mouse scRNA-seq data

After spatial inference of hepatocytes, the spatial expression profiles were further characterized by a model selection approach. Genes with > 300 total reads in at least one condition were retained. For these 6235 genes, we employed the ’glmmTMB’ R package ^78^ to fit a generalized linear mixed-effects (GLMM) model (with a log link function), accounting for negative binomial noise and utilizing the Broyden–Fletcher–Goldfarb–Shanno optimization method. To address numeric instability for genes not expressed in at least one condition (less than 10 total counts), we performed an imputation using random draws from a negative binomial distribution with mean *μ* = *e*^-4^ and dispersion 0.1. Spatial coordinates across all samples were normalized to the range of 0 to 1 for consistent comparisons. Ten models, with incrementally increased complexity, were considered for each gene. These models included fixed effects, with intercepts and slopes being either sex, or time-specific, along with possible interaction terms, and random effects for the intercept and slope in each animal. The library size, computed by summing all gene counts except mitochondrial genes, served as an offset in the GLMM. A Bayesian Information Criterion (BIC) was employed, assigning genes to the model exhibiting the lowest BIC. To optimize computational efficiency, we parallelized the analysis at the gene level using the ’mclapply’ function from the parallel R package. We kept the estimated fixed effect coefficients from the best-fitting model for each gene. Note that the current simplest model with linear dependency in the positions was not designed to capture genes with peaks in the midlobular region, of which previous analysis identified only very few ^26^. Moreover, the assignment to complex models with interacting effects of sex and time on the shape of zonation profiles did not yield many genes (15 genes in models 9 and 10, BIC, S. Table 3), and for those models the assignments seem sensitive to noise. Therefore we did not consider those genes further.

### Fenofibrate injections

Fenofibrate was injected intraperitoneally at 80 mg per kg body mass for 5 straight days, at ZT0. Due to its low solubility and instability in water, fenofibrate was dissolved in DMSO in 10x working concentration and frozen at -20 °C. Upon thawing of the solution, medical grade 0.9% NaCl was added to the final volume, resulting in a suspension (rather than solution), which was shaken before loading into the syringe and injecting. Tissue was isolated at two time points that followed the 5th injection, ZT10 and ZT22.

### Histology

#### Collection and freezing of mouse tissue

Tissue for histology was isolated, placed into cryo molds, covered with 2% carboxymethylcellulose (CMC) in distilled water (prepared by sonication of CMC in water), and placed into a deep section of dry ice, allowing for even and quick freezing. Alternatively, the tissue was frozen without CMC in the same manner. The tissue was subsequently cryosectioned at 10 µm, placed on glass slides and stored at -80 °C until processing.

#### Immunofluorescence of global zonation markers

Triple immunofluorescence was performed manually on mouse liver 10 µm fresh frozen sections. Tissues were fixed with 4% PFA for 15 min at RT before permeabilization with 0.1% Tween in 1x PBS for 10 min. After 30 min of blocking with 1% BSA in 1x PBS, primary antibodies were incubated simultaneously overnight at 4 °C: rabbit anti-CYP2E1 (Abcam ab28146, 1 : 300), mouse anti-GS/GLUL (Santa Cruz sc-74430, clone E-4, 1 : 100) and goat anti-E-CAD (R&D systems AF478, 1 : 200). After washes with 1x PBS, secondary antibodies were applied simultaneously for 40 min at RT: donkey anti-rabbit Alexa568 (Thermo Fisher A10042, 1 : 1000), donkey anti-mouse Alexa647 (Thermo Fisher A31571, 1 : 1000) and donkey anti-goat Alexa488 (Thermo Fisher A11055, 1 : 1000). Sections were counterstained with DAPI and mounted with FluoromountG (Bioconcept 0100-01).

#### VLDLR immunohistochemistry (mouse sections)

The immunohistochemistry was performed using the fully automated Ventana Discovery ULTRA (Roche Diagnostics, Rotkreuz, Switzerland). All steps were performed on the equipment with Ventana solutions except if mentioned. Briefly, 10 µm mouse liver fresh frozen sections were fixed with 4% PFA for 15 min at RT, washed with 1x PBS before loading wet on the automated device. The primary antibody, goat anti-VLDLR (R&D Systems AF2258, diluted 1 : 50 in 1% BSA in 1x PBS) was incubated 1 h at 37 °C. After incubation with goat Immpress HRP (Ready to use, Vector Laboratories), chromogenic revelation was performed with ChromoMap DAB kit. Sections were counterstained with Harris hematoxylin and permanently mounted.

#### VLDLR immunohistochemistry (human sections)

Human tissue was processed using the Ventana Discovery ULTRA machine with Ventana solutions except if mentioned. Dewaxed and rehydrated 4 µm paraffin sections were pretreated with heat using the CC1 solution for 40 minutes at 95 °C. For DAB staining, the primary antibody, goat anti-VLDLR (R&D Systems AF2258), was used in a dilution of 1 : 25 (1% BSA in 1x PBS), 1 h incubation at 37 °C. After incubation with goat Immpress HRP (Ready to use, Vector Laboratories), chromogenic revelation was performed either with the ChromoMap DAB kit or with the DISCOVERY Purple Kit. Sections were counterstained with Harris hematoxylin and permanently mounted. Given that the VLDLR antibody is not validated for clinical use, we further assessed its specificity in a western blot and in a control tissue, the pancreas (SFig. 25). Given a higher background stain in the two needle biopsies (different tissue fixation), these were excluded from the quantification.

#### Staining of neutral lipids

The staining of neutral lipids was done with BODIPY 493/503 (Cayman Chemical). Flash frozen tissue, cut at 10 µm, was thawed and incubated with BODIPY in 1x PBS for 1 h at room temperature under gentle shaking, protected from light. After incubation, the tissue was washed three times with 1x PBS for 3 min. Sections were counterstained with DAPI, which was added to the first PBS wash. After the final wash, the samples were dried, protected from light, and immediately imaged without mounting.

#### Imaging

Imaging was done with the Olympus VS120 and Olympus VS200 slide scanners and Leica DM5500 widefield microscope. Images were subsequently analyzed using QuPath ^79^ and FIJI software suites.

#### Coronary artery

The evaluation of atherosclerosis was performed on coronary artery H&E images from GTEx. For each donor, the coronary artery sections were assessed by a clinical vascular pathologist (M. R.) and cross-referenced to the metadata and bulk tissue expression, as described above. For the group of individuals ≤ 49 years of age, 2 samples were excluded based on their medical history (procedures in the area of interest which could affect the results). When assessing the donors of all age groups, no individuals were excluded, as the large sample size prevented individual donors from skewing the data. The prevalence of a condition was calculated cumulatively in function of age (what percentage of individuals below a given age show the condition), with a subsequent linear fit across the data-points for easier visualization.

### Electron microscopy (EM)

Two different preparation methods were used to image VLDL particles in hepatocytes with electron microscopy. A first, optimized for visualizing cell ultrastructure at high resolution with transmission electron microscopy, appears to leave the VLDL particles unstained, therefore light gray. While the second method, optimized for imaging cell ultrastructure in serial images with block face scanning electron microscopy, renders the particles heavily stained, therefore black (SFig. 13). This dark staining of these particles is similar to that seen in previous methods where the tissue is stained using osmium tetroxide as the primary fixative ^53^.

#### Mouse perfusion

After euthanasia, and immediately after the heart had stopped beating, adult mice were perfused with a buffered mix of 1.2% glutaraldehyde and 2.0% paraformaldehyde in 0.1 M phosphate buffer (pH 7.4). The livers were removed 2 hours after the perfusion had finished, and were then vibratome sectioned at 100 µm thickness.

#### Preparation of liver tissue for transmission electron microscopy

These sections were then washed with cacodylate buffer (0.1 M, pH 7.4), postfixed for 40 min in 1.0% osmium tetroxide with 1.5% potassium ferrocyanide, and then 40 min in 1.0% osmium tetroxide alone. They were then stained for 30 min in 1% uranyl acetate in distilled water before being dehydrated through increasing concentrations of ethanol and then embedded in Durcupan ACM (Fluka, Switzerland) resin. The slices were finally mounted between two glass slides and placed in an oven for 24 hours at 65 °C. Once the resin had hardened, a region of interest from a single slice was cut away from the rest and glued to a blank resin block, with cyanoacrylate glue, and thin (50 nm thick) serial sections were cut with a diamond knife (Diatome, Switzerland). These were collected onto pioloform support films on single slot copper grids and contrasted with lead citrate and uranyl acetate. Images were taken with a transmission electron microscope at 80 kV (Tecnai Spirit, FEI Company with Eagle CCD camera).

#### Preparation of liver tissue for serial block face scanning electron microscopy

The vibratome slices of fixed liver tissue were postfixed in potassium ferrocyanide (1.5%) and osmium (2%), then stained with thiocarbohydrazide (1%) followed by osmium tetroxide (2%) alone. They were finally stained overnight in uranyl acetate (1%), washed in distilled water at 50 °C before being stained with lead aspartate at pH 5.5 at the same temperature. The slices were then dehydrated in increasing concentrations of alcohol and embedded in Durcupan resin and hardened at 65 °C for 24 h. Once hardened, the region of interest was cut away from the rest of the slice with a scalpel blade. This piece was glued to an aluminum holder with conductive cement and then placed inside the scanning electron microscope (Merlin, Zeiss NTS) integrated with an in-chamber ultramicrotome device (3View, Gatan). The ultramicrotome was set to remove 50 nm thick slices from the surface of the block and images of the region were taken after each removal. An imaging voltage of 1.7 kV was used and a pixel size of 7 nm. Backscattered electrons were collected with a Gatan backscattered electron detector with a pixel dwell time of 1 µs.

### Tracer studies

129S6/SvEv (Taconic) mice, aged 4-6 months, were placed under an IR lamp 5 min before tail vein injections to ensure vasodilation and visualization of the vein. The mice were then placed into a restrainer, and injected with 200 µl of the buffered solution of VLDL particles containing the DiI (1,1’-dioctadecyl-3,3,3’3’-tetramethylindocarbocyanine perchlorate) fluorophore (Kalen Biomedical). After 15 min, the mice were injected with 150 mg per kg body mass pentobarbital, and dissected 5 min later. At that time-point, all reflexes were absent in all of the mice, and death was confirmed with decapitation. Blood was removed from the liver by perfusion with PBS via the spleen, whereas other imaged organs were cut open and flushed of blood in PBS. Organs were collected, immediately imaged in the IVIS in vivo imaging system (PerkinElmer), in combination with the Living Image 4.5 software.

