## Supplemental data for "A sexually dimorphic hepatic cycle of periportal VLDL generation and subsequent pericentral VLDLR-mediated lipoprotein re-uptake"

### *Extended data*

#### *Tables*

*Supplementary Table 1:* ***List of pathology associated genes and literature sources.*** *The genes in the table were shown to be either up- or downregulated in the associated conditions, or were shown to have a causal relationship with the pathology. The different capitalization refers to either rodent or human genes, or proteins if not italicized.*

| **Condition** | **Tissue or cell type** | **Genes** | **Reference** |
|---|---|---|---|
| NAFLD and NASH | liver bulk | *IL17RA* | Giles et al., 2015, PMID: 26028039 |
| NAFLD and NASH | liver bulk | *GCKR* | Pirola et al., 2018, PMID: 30202818 |
| NAFLD and NASH | liver bulk | *PNPLA3*, *TMSF6*, *GCKR*, *HSD17BB13*, *MARC1*, *PPP1R3B*, *LEPR*, *APOE*, *GPAM*, *MBOAT7*, *LYPLAL1*, *PYGO1* | Du et al., 2021, PMID: 34950105 |
| NAFLD and NASH | liver bulk | *Pnpla3* | Smagris et al., 2015, PMID: 24917523 |
| HCC | liver bulk | *SH2D4A* | Quagliata et al., 2014, PMID: 24315626 |
| HCC | liver bulk | *SH2D4A*, *SORBS3*, *IL6* | Ploeger et al., 2016, PMID: 27311882 |
| HCC | liver bulk | *CHD1L*, *CKS1B*, *JTB*, *SHC1*, *BCL9*, *ARNT*, *TPM3*, *MUC1*, *NTRK1*, *CD1D*, *MPZL1*, *MYC*, *DDEF1*, *MLZE*, *HEY1*, *CTHRC1*, *BOP1*, *SGCE*, *DYNC1I1*, *PEG10*, *ING2*, *ADH3*, *ADH1C*, *ADH1A*, *ADH6*, *DLC1*, *CCDC25*, *ELP3*, *PROSC*, *SH2D4A*, *SORBS3*, *MCPH1*, *KIAA1456*, *TUSC3*, *ZDHHC2*, *ARD1B*, *SEPT11*, *M6P*/*IGF2R* | Niu et al., 2016, PMID: 27895396 |
| HCC | liver bulk | genes encoding ADH and ALDH enzymes | Meroni et al., 2018, PMID: 30513996 |
| HCC | liver bulk | *Dpyd* | Zhu et al., 2018, PMID: 29358721 |
| HCC | liver bulk | *Gna14* | Song et al., 2022, PMID: 33500727 |
| HCC, fibrosis, steatosis | liver bulk | *Pdgfc* | Campbell et al., 2005, PMID: 15728360 |
| HCC, fibrosis | hepatocytes | *Pdgfc* | Jin Lim et al., 2018, PMID: 30509307 |
| liver microenvironment of hepatic metastases | liver bulk | *Hgf*, *Il8*, *Tgfb1*, *Mmp2*, *Postn*, *Mmp9*, family members encoding NFkB, peptidyl arginine deiminases, collagen 1 and 4, CEA, MAPK, VEGF | Williamson et al., 2019, PMID: 31263976 |
| alcoholic liver disease | liver bulk | *PNPLA3*, *TM6SF2*, *MBOAT7*, *SOD2*, *MMP3*, *TGFB1* | Meroni et al., 2018, PMID: 30513996 |
| alcoholic liver disease | liver bulk | genes encoding ADH and ALDH enzymes | Meroni et al., 2018, PMID: 30513996 |
| DILI | liver bulk | genes encoding SLC, OATP, OAT, OCT family members, *SLCO1B1*, *SLCO1B3*, *SLCO2B1*, *ABCB11*, *ABCG2*, *ABCB4*, *CYP2E1*, *PTPN22* | Andrade et al., 2019, PMID: 31439850 |
| DILI | liver bulk | *NAT2* | Zhang et al., 2018, PMID: 30047605 |
| DILI | liver bulk | *IL1B* | Bonkovsky et al., 2018, PMID: 30359443 |
| DILI | liver bulk | *SOD2*, *GPX1* | Lucena et al., 2010, PMID: 20578157 |
| monogenic diseases | liver bulk | *AGXT*, *ARG1*, *CPS2*, *ASS1*, *ASL*, *OTC*, *GLUL* | Martini et al., 2023, PMID: 36693201 |
| monogenic diseases | liver bulk | *JAG1*, *NAGS*, *ATP7B*, *ATP8B1*, *FAH*, *CFH*, *TTR*, *UGT1A1* | Fagiuoli et al., 2012, PMID: 23578885 |
| cirrhosis | $CD74^+$ cells | *FYN, IFNG, KLRD1, HLA-G, ID2, ETS1, IRF1, PRDM, CD8A, CCL4, CCL3, CXCR4, JUN, CCL5, SOCS3, IRF8, NR4A2, IKZF3, REL* | Liu et al., 2022, PMID: 35903097 |
| fibrosis | stellate cells | *MT1* (also *MT1A*) | Xu et al., 2013, PMID: 23620539 |

| | | | |
|---|---|---|---|
| fibrosis | liver bulk | *Hsd17b13* | Luukkonen et al., 2023, PMID: 36669104 |
| fibrosis | stellate cells | *AHSG* | Yilmaz et al., 2010, PMID: 20926473 |
| fibrosis | stellate cells | *ZBTB16* | Gerhard et al., 2020, PMID: 32258441 |
| inflammation | $CD74^+$ cells | *Gm26917* | Zhao et al., 2022, PMID: 36386180 |
| cardiovascular disease | / | *PNPLA3*, *TM6SF2* | Brouwers et al., 2019, PMID: 31713012 |
| cardiovascular disease | / | *PNPLA3*, *TM6SF2* | Meroni et al., 2018, PMID: 30513996 |
| cardiovascular disease and lipid homeostasis | liver bulk | *PNPLA3* | Gluchowski et al., 2017, PMID: 28428634 |
| cardiovascular disease, NAFLD, fibrosis, HCC | / | *TM6SF2* | Luo et al., 2022, PMID: 34532996 |
| plasma TAG levels and NAFLD | / | *TM6SF2* | Mahdessian et al., 2013, PMID: 24927523 |
| atherosclerosis | serum (produced by the liver) | *SAA1*, *SAA2*, *Saa3* | Shridas and Tannock, 2019, PMID: 31135596 |
| atherosclerosis | serum (produced by the liver) | *Saa3* | Thompson et al., 2018, PMID: 29175652 |
| cardiovascular disease | liver bulk | *APOA5*, *APOB*, *APOC3*, *ABCG5/8*, *SORT1*, *LDLR*, *HNF1A*, *LPA*, *LPL*, *PCSK9*, *ABO*, *LRP1*, *ANGPTL4*, *TRIB1*, *SCARB1*, *PLTP* | Bjorkegren and Lusis, 2022, PMID: 35504280 |

*Supplementary Table 2:* ***Gene expression in hepatocytes in function of sex, time and lobular layer.*** *For each cell (rows) in the hepatocyte subcluster with ≥ 3% mitochondrial reads, the gene (columns) expression is given in raw counts. The sex, time and the lobular layer of each cell are also provided (columns). The layers were calculated by binning the spatial coordinates into 12 bins, 1 corresponds to the most pericentral and 12 to the most periportal layer. Please refer to the corresponding .csv file.*

*Supplementary Table 3:* ***Model selection.*** *Effects of sex, time and lobular layer on the expression profiles of all of the genes were statistically assessed by a model selection approach (BIC - Bayesian Information Criterion; Methods). Models 1-5 describe non-zonated gene expression profiles. Model 1 encompasses non-zonated genes which have a constant mean across the four different groups (female ZT10, female ZT22, male ZT10, male ZT22). Genes in model 2 are non-zonated with an effect of sex on the mean, and model 3 genes are non-zonated with an effect of time on the mean. Model 4 genes exhibit an effect of sex at one time-point or an effect of time in one sex (three different expression means). Model 5 genes are non-zonated and have independent means in all four groups. Models 6-10 describe zonated genes. Genes in model 6 have the same slope across conditions, genes in model 7 show an effect of sex on the slope of zonation, and genes in model 8 show an effect of time on zonation. Genes in model 9 have an interactive effect of sex and time on the slope, and genes in model 10 are those in which each condition (female ZT10, female ZT22, male ZT10, male ZT22) has its own slope. Please refer to the corresponding .csv file.*

*Supplementary Table 4:* ***Pathological assessment of coronary artery specimens.*** *All coronary artery images of GTEx donors, for which there is corresponding hepatic bulk RNA-seq data, were assessed by a clinical cardiovascular pathologist. Publicly accessible donor data and the assessment are shown in the table. The images can be accessed via the GTEx Portal (GTEx Histology Viewer). Please refer to the corresponding .csv file.*

*Supplementary Table 5:* ***Clinical characteristics of human tissue and donors.*** *For each donor, we provide the age, body mass index (BMI), information on lipid lowering medication, the reason for resection and the assessment of the tissue by the pathologist. The pathologist did not have access to the donors' medical history or reason for resection to enable a blind and unbiased assessment of the sample. A BMI over 25 (overweight) is highlighted, the asterisk (*) marks samples where the body weight and height for BMI calculation were not recorded around the period of resection. In samples that came from cancer resections, a section from the healthy surrounding tissue was used for VLDLR immunostaining.*

| # | Sex | Age | BMI | Lipid lowering medication | Reason for resection | Unbiased pathological assessment of tissue used for staining, without clinical information |
|---|---|---|---|---|---|---|
| 1 | female | 25 | 24.4 | not treated with ATC C10 | hepatocellular carcinoma | mild portal inflammation |
| 2 | female | 27 | NA | not treated with ATC C10 | hemihepatectomy due to nodular hyperplasia | normal |
| 3 | female | 36 | **26.8** | C10AA05 | hemihepatectomy due to cysts | dense portal infiltrates, no tumor, no steatosis |
| 4 | female | 39 | NA | not treated with ATC C10 | cholecystitis | no tumor in the section, but likely somewhere else in the tissue due to high neutrophil count, mild but clinically insignificant steatosis, no portal inflammation, mild lobular inflammation |
| 5 | female | 42 | 20.9 | not treated with ATC C10 | echinococcus | portal and lobular inflammation, suspected echinococcus or sclerosing hemangioma, no steatosis, localized fibrotic tissue |
| 6 | female | 46 | **28.2** | not treated with ATC C10 | hepatocellular adenoma | fibrosis around bile ducts, no significant steatosis, mild portal and lobular infiltrates, suspected primary sclerosing cholangitis, cholangiocellular carcinoma |
| 7 | female | 46 | 21.5 | not treated with ATC C10 | intestinal adenocarcinoma | adenocarcinoma (necrotic), prominent lipofuscin depositions in adjacent liver tissue, no steatosis, no significant portal or lobular inflammation |
| 8 | male | 45 | 19.9 | not treated with ATC C10 | Crohn's disease (biopsy) | normal |
| 9 | male | 40 | 22.8 | C10AA07 | cavernous hemangioma | hemangioma or angiosarcoma, proliferation of blood vessels, limited liver tissue, portal inflammation, no steatosis, not possible to assess lobular inflammation due to amount of tissue |
| 10 | male | 30 | **29.0** | not treated with ATC C10 | hepatocellular carcinoma | suspected HCC, portal inflammation, mild lobular inflammation, no steatosis |
| 11 | male | 37 | 24.2 * | not treated with ATC C10 | suspected steatosis (biopsy) | normal |
| 12 | male | 26 | **27.1** | not treated with ATC C10 | echinococcus | echinococcus, extensive lipofuscin deposition, no steatosis, no significant inflammation except next to the parasite, no tumor |
| 13 | male | 49 | 22.7 | not treated with ATC C10 | hepatic adenoma | normal |
| 14 | male | 45 | **28.8** | not treated with ATC C10 | hepatocellular carcinoma | pronounced ductular reaction, no steatosis, no lobular inflammation, mild portal inflammation, no tumor |
| 15 | male | 35 | 20.6 | not treated with ATC C10 | intestinal adenocarcinoma | giant cells bordering sclerosing carcinoma or echinococcus, portal inflammation, considerable lipofuscin depositions, no lobular inflammation, no steatosis |
| 16 | male | 46 | 24.7 | not treated with ATC C10 | intestinal adenocarcinoma | no H&E available |
| 17 | female | 64 | 21.1 | not treated with ATC C10 | adenocarcinoma | localized presence of giant cells and traces of previous surgery, no significant steatosis, no |

| | | | | | | |
|---|---|---|---|---|---|---|
| | | | | | | significant lobular inflammation, mild to moderate portal inflammation |
| 18 | female | 57 | 23.3 | not treated with ATC C10 | adenocarcinoma | no tumor, no steatosis, focal portal inflammation, no lobular inflammation |
| 19 | female | 69 | **27.9** | not treated with ATC C10 | echinococcus and cavernous hemangioma | normal |
| 20 | female | 60 | 21.2 | not treated with ATC C10 | biliary cysts | mild ductular reaction |
| 21 | female | 77 | 18.0 * | not treated with ATC C10 | intestinal adenocarcinoma | portal inflammation, tumor in blood vessel (suspected adenocarcinoma), lipofuscin deposition |
| 22 | female | 79 | 22.5 | C10AA07 | intestinal adenocarcinoma | normal |
| 23 | female | 69 | **25.9** | C10AA05 | adenocarcinoma | dense portal infiltrates, lobular inflammation, very pronounced lipofuscin deposits, no steatosis, no tumor |
| 24 | female | 69 | **29.5** | not treated with ATC C10 | cholangiocellular carcinoma | probable HCC, bordering tissue with dense portal infiltrates, lobular inflammation, no steatosis |
| 25 | female | 79 | 24.8 | not treated with ATC C10 | adenocarcinoma | normal, minor sample artifacts visible |
| 26 | male | 80 | **31.1** | C10AA05 | hepatocellular carcinoma | no tumor, no significant steatosis, mild portal inflammation, no lobular inflammation |
| 27 | male | 67 | NA | not treated with ATC C10 | melanoma | ductular reaction (suspected tumor in a different region of the liver), mild portal inflammation, no lobular inflammation, no steatosis |
| 28 | male | 70 | 21.5 | C10AA07 | neuroendocrine tumor | mild portal inflammation |
| 29 | male | 57 | 23.8 | not treated with ATC C10 | neuroendocrine tumor | non-significant steatosis, no inflammation, no tumor |
| 30 | male | 61 | 22.5 | not treated with ATC C10 | colorectal carcinoma | adenocarcinoma contamination, lipofuscin depositions, mild portal inflammation, mixed micro- and macrovesicular steatosis (5%-10%) |
| 31 | male | 65 | **26.3** | not treated with ATC C10 | rectal adenocarcinoma | normal |
| 32 | male | 65 | 21.2 | not treated with ATC C10 | intestinal adenocarcinoma | mild microvesicular steatosis (5%-10%), portal inflammation, no tumor, no lobular inflammation, very pronounced lipofuscin depositions |
| 33 | male | 54 | **25.5** | not treated with ATC C10 | neuroendocrine tumor | large tumor, unable to determine type of carcinoma; in adjacent liver tissue no steatosis, portal and lobular inflammation |

#### *Figures*

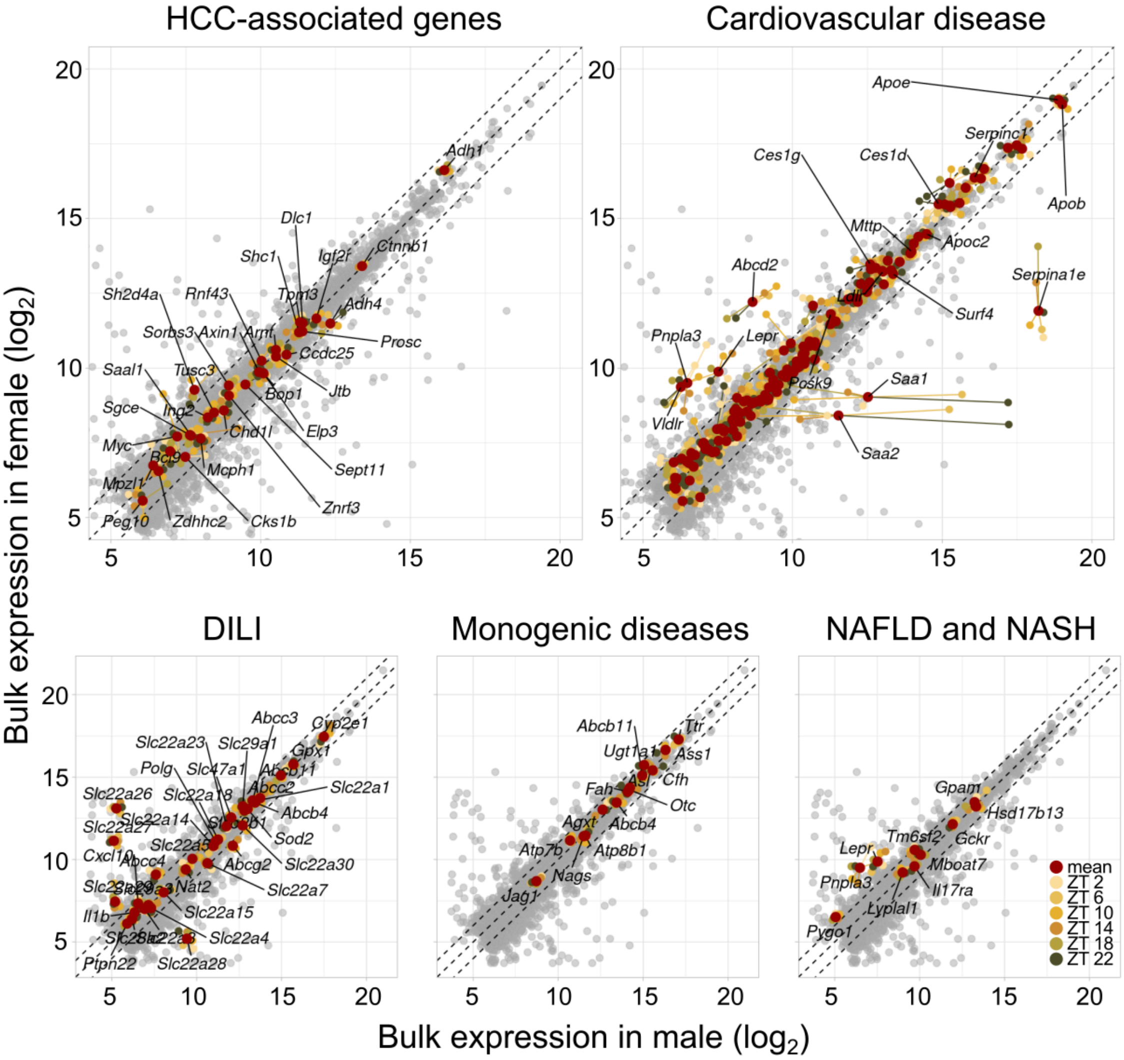


*SFig. 1:* ***Effects of sex and time on pathology-associated genes in bulk RNA-seq (reanalysis of data from ref. 7).*** *Hepatic expression of genes that are either involved in liver pathologies or that are mainly expressed in the liver and encode secreted proteins that affect systemic diseases. The male expression is plotted on the x-axis, while the female expression is plotted on the y-axis. For genes of interest, their mean expression, as well as their expression at different time points is plotted, as per legend. The tissue isolation was performed as a time-series on temporally entrained animals. Bulk sequencing in C56BL/6 mice reveals that few pathology-associated genes show a 2-fold difference (dashed lines) in expression between females and males. In HCC-associated genes, Sh2d4a, commonly downregulated in HCC, shows higher female expression. Adh1 is the highest expressed alcohol dehydrogenase, showing a small female bias, whereas Adh4 is higher in males. Some genes, such as Myc, Cks1b, Mpzl1, Sgce and Peg10, show 2-fold or higher daily expression amplitudes. The highest sexual dimorphism is seen in cardiovascular disease-associated genes, with an 8-fold female to male expression ratio of Vldlr. There are also considerable differences in the coagulation cascade and inflammatory processes, serum amyloid A 1, 2 and 3 as well as in antithrombin (Serpinc1) and alpha-1 antitrypsin (Serpina1). At the same time, the low density lipoprotein receptor (Ldlr) and apolipoproteins (Apos) are comparable between males and females. Effects of sex on monogenic liver disease genes are negligible, with the exception of Ugt1a1. Contrary, there is considerable sexual dimorphism in solute carriers, which are commonly associated with drug-induced liver injury (DILI). Some results presented in this figure were extrapolated from*

*human data onto mouse bulk RNA-seq. Therefore, interpretation should be done with caution, considering potential differences in hepatic metabolism and circadian rhythm between species (n = 12, per sex; n = 2, per time-point).*

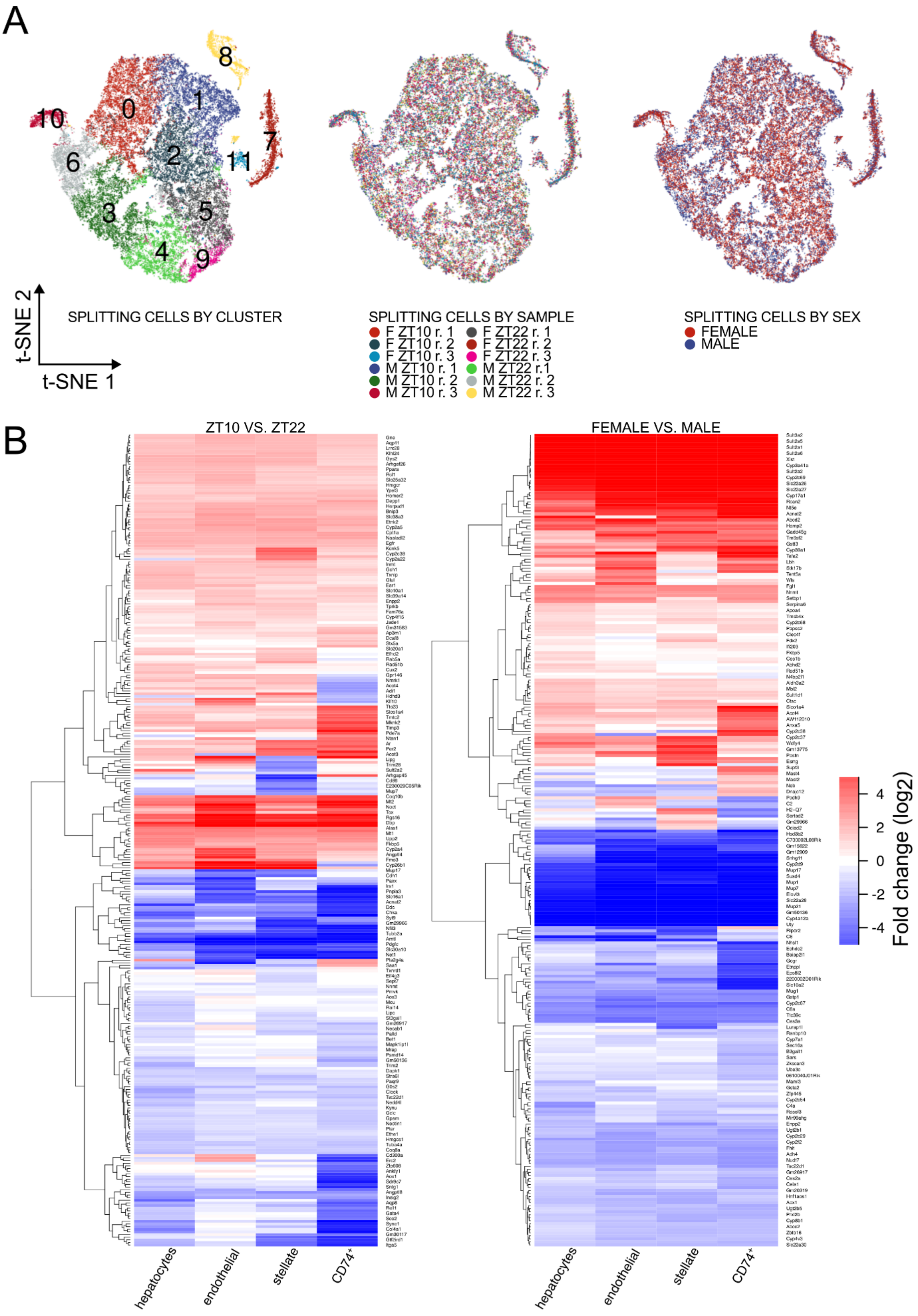


SFig. 2: ***scRNA-seq sample integration and clustering.*** *(A) After integration, the samples do not separate by sample ID or sex in the t-SNE plots, allowing for the identification of different cell types. All cell types appear in all samples and there are no individual clusters that are sample specific (n = 12). (B) We assessed effects of time*

*(left panel, ZT10 vs. ZT22; positive fold changes denote genes that were more highly expressed at ZT10) or sex (right panel, female vs. male; positive fold changes denote genes that were more highly expressed in females). The set of genes was selected by the criterion that the $log_2$ fold change needs to be at least 1 in at least one cell type. Effects of time and sex are largely conserved between cell types, with the exception of specific modules of temporal and sex-dimorphic regulation in a subset of genes in $CD74^+$ and stellate cells. Every second gene/line is named.*

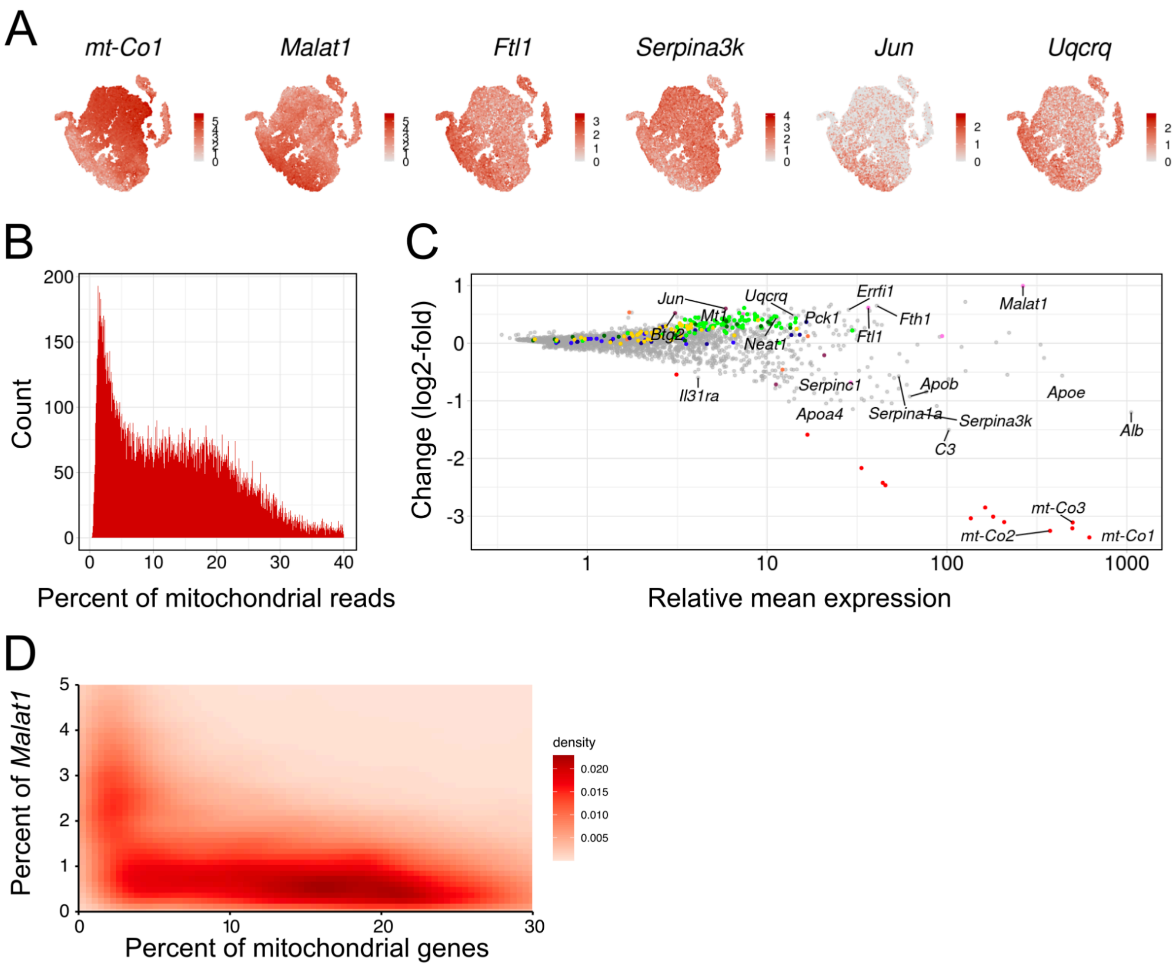


*SFig. 3:* ***scRNA-seq reveals a mt-Co1-negative subcluster of hepatocytes.*** *(A) t-SNE plots reveal that the main cluster of hepatocytes includes a subset of cells (lower left of the main cluster in the t-SNE) that have low mt-Co1 and Serpina3k expression, but higher expression of Malat1 (RNA processing), Ftl1 (iron homeostasis), Jun (proliferation, differentiation, apoptosis, stress response) and Uqcrq (mitochondrial respiratory chain). (B) Hepatocytes exhibit a bimodal distribution of mitochondrial read percentage. To split the hepatocytes into those with low and high mitochondrial fractions, a cutoff of 3% mitochondrial genes was chosen. (C) The cutoff allows for the separation of the mt-Co1-negative hepatocytes. The change vs. mean expression plot shows how the mitochondrial gene depleted cluster (mitochondrial fraction < 3% compared to all genes) differs from other (≥ 3%) hepatocytes (y-axis) in function of average expression of a gene across both the low and high mitochondrial fraction hepatocytes (x-axis). In the low mitochondrial group, we confirmed a downregulation of mitochondrial genes (red) and genes encoding secreted proteins, such as serpins and apolipoproteins. Contrary, these cells show an upregulation of ribosomal proteins (bright green), glucose metabolism (blue), and especially gluconeogenesis-associated genes (dark blue), Nduf genes (yellow), cytochrome o oxidases (dark green), iron metabolism-associated genes (pink), angiogenesis (orange), and stem cell and proliferation/malignancy markers (plum). One surprising observation is the upregulation of Nduf and Cox genes, and Uqcrq, which are, among other cellular functions, involved in the mitochondrial transport chain. (D) A density plot in which cells split into two clusters, one with low mitochondrial reads and high Malat1, and one with low Malat1 and higher mitochondrial reads (n = 12).*

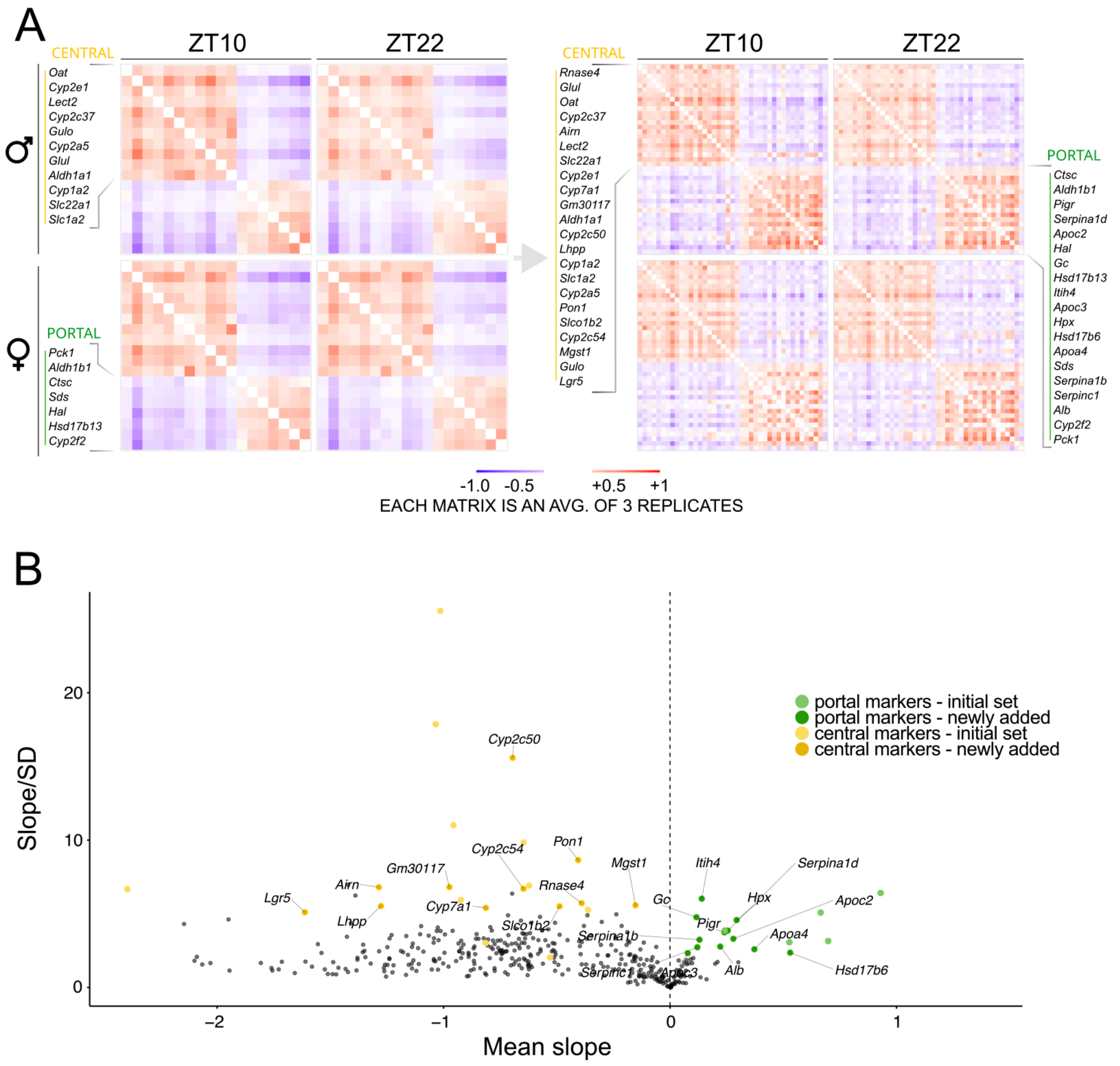


*SFig. 4: **Spatial reconstruction of hepatocytes with a mitochondrial fraction of ≥ 3%.** (A) A Pearson's correlation analysis reveals that the correlation between the selected portally and centrally biased genes is conserved independently of time and sex. The initial set of zonation markers (left panel) and the extended set of zonation markers (right panel) are shown (n = 12). (B) These additional zonation markers were identified with the model for inference of spatial position. For each candidate gene and for each mouse, the slope of the expression (over the central to portal axis) was calculated. Subsequently, the mean and standard deviation across mice was calculated for each gene and then plotted. The absolute value of the slope's mean to standard deviation ratio (signal to noise) was plotted against the slope's mean. The yellow dots represent central and green portal genes, and fainted dots the initial set of marker genes. Newly detected markers are named (n = 11).*

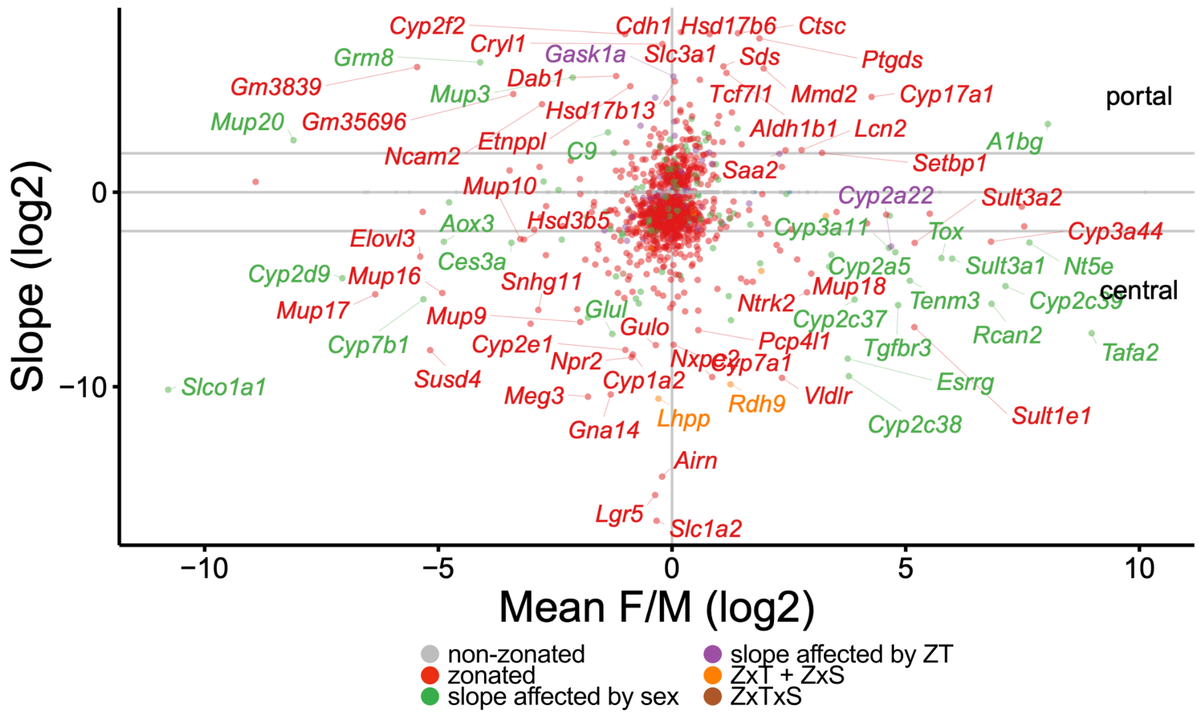


*SFig. 5:* ***Zonation of sexually dimorphic genes.*** *Zonation is represented by the average slope (negative slopes characterize central profiles) across all conditions (y-axis) and compared to the female to male expression ratio (x-axis). Named genes are highly zonated (a 4-fold or higher difference between the portal to central expression or vice versa). Genes in the corners are additionally highly sexually dimorphic, including cytochromes, sulfotransferases, solute carriers and Vldlr. Slopes were calculated by averaging, across conditions, fixed effect coefficient estimates from the best fitting model based on the BIC. The different coloured points represent different models based on the slope estimates from the model selection approach.*

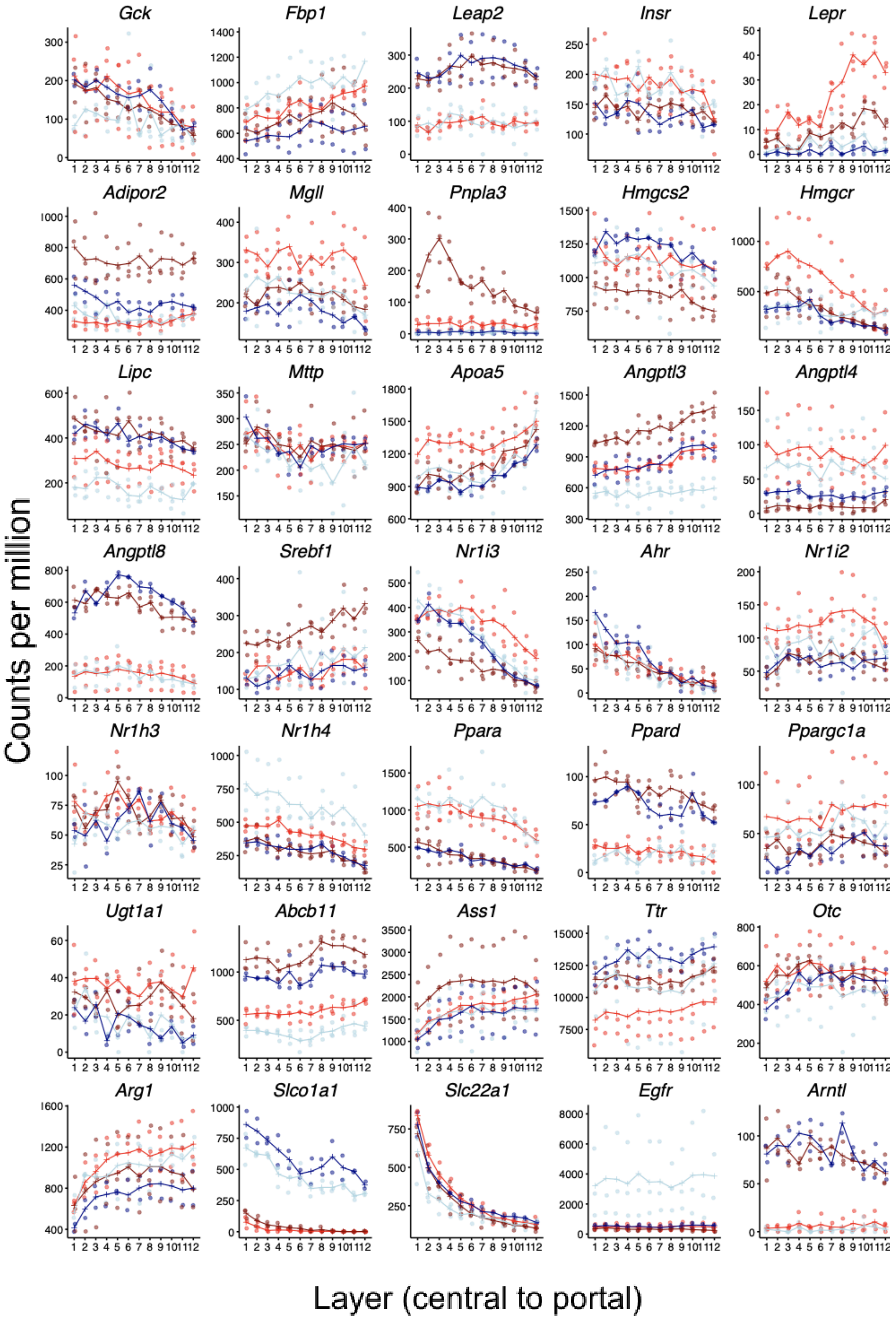
Gck
Fbp1
Leap2
Insr
Lepr
Adipor2
Mgll
Pnpla3
Hmgcs2
Hmgcr
Lipc
Mttp
Apoa5
Angptl3
Angptl4
Angptl8
Srebf1
Nr1i3
Ahr
Nr1i2
Nr1h3
Nr1h4
Ppara
Ppard
Ppargc1a
Ugt1a1
Abcb11
Ass1
Ttr
Otc
Arg1
Slco1a1
Slc22a1
Egfr
Arntl
Counts per million
Layer (central to portal)

*SFig. 6:* ***Expression of metabolically relevant and pathology associated genes in function of time, sex and lobular layer.*** *Expression of metabolically relevant and pathology associated genes, shown as normalized counts per million (y), excluding mitochondrial genes for normalization, in function of central to portal position (x), binned into 12 layers in relation to the central vein (n = 11). Glucokinase (Gck) shows a pericentral bias. The fructose-bisphosphatase 1 (Fbp1) has higher periportal expression, and is affected by time in males, with higher expression at ZT10. The ghrelin receptor antagonist Leap2 is affected by time, with higher expression at ZT22, in the fed state. Whereas the insulin receptor does not seem to be affected by sex, the leptin receptor shows striking sexual dimorphism and highest expression in females at ZT10 periportally. The adiponectin receptor 2 (Adipor2) is not affected by space, but shows higher expression in the female ZT22 group compared to other conditions. The monoglyceride lipase (Mgll) shows higher expression at ZT10. Pnpla3 shows a pericentral expression in the female ZT22 group. Hmgcs2, crucial for ketone body synthesis, shows higher expression in the fasted state. The pericentral expression of Hmgcr suggests a pericentrally biased cholesterol synthesis. The hepatic lipoprotein lipase (Lipc) shows higher expression at ZT22, in the fed state. Mttp does not show major effects of time, sex or space. Apolipoprotein A5 is higher periportally. Angiopoietin-like protein coding genes exhibit predominantly effects of time and in the case of Angptl3 also sexual dimorphism. There is higher female expression of Srebf1 at ZT22, encoding the sterol regulatory element-binding protein 1, a master regulator of lipid metabolism. The xenobiotic sensor Car/Nr1i3, responsible for transcriptional regulation of cytochromes, shows a higher pericentral expression. Ppara is higher in anticipation of feeding, and Ppard shows higher expression in the fed state. The monogenic disease associated gene expression profiles reveal a higher Ugt1a1 expression in females, effects of time on the ATP binding cassette transporter Abcb11, and a periportal bias for Ass1 and Arg1. Of note, the sexual dimorphism in Ugt1a1 could be causal for sex differences in bilirubin. Slco1a1, encoding OATP1A1, is sexually dimorphic, with higher male expression. Slc22a1, encoding OCT1, is pericentral. The epidermal growth factor receptor is increased in males at ZT10. The molecular clock gene Arntl/Bmal1 peaks at the end of the feeding phase.*

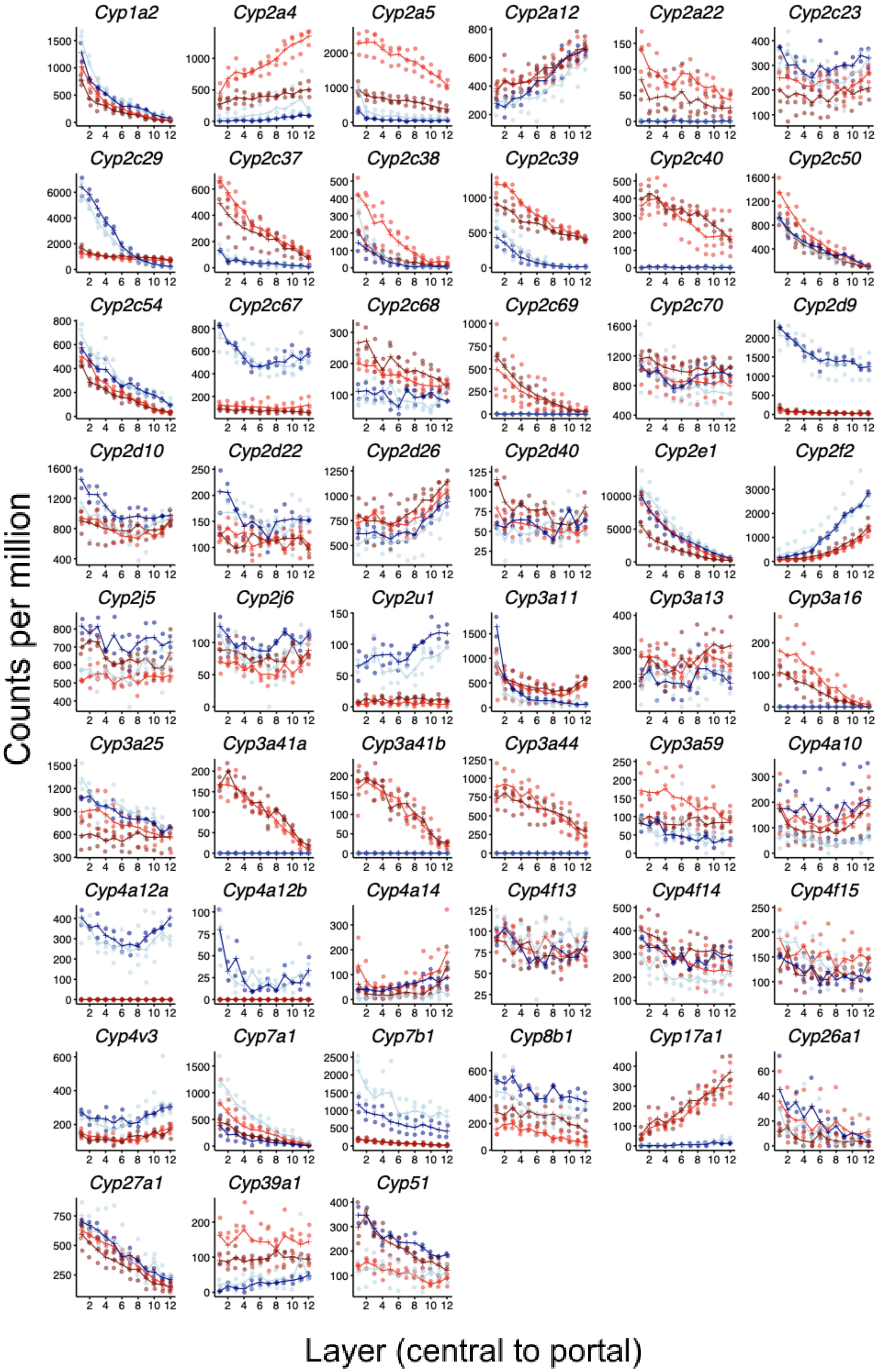
Cyp1a2
Cyp2a4
Cyp2a5
Cyp2a12
Cyp2a22
Cyp2c23
Cyp2c29
Cyp2c37
Cyp2c38
Cyp2c39
Cyp2c40
Cyp2c50
Cyp2c54
Cyp2c67
Cyp2c68
Cyp2c69
Cyp2c70
Cyp2d9
Cyp2d10
Cyp2d22
Cyp2d26
Cyp2d40
Cyp2e1
Cyp2f2
Cyp2j5
Cyp2j6
Cyp2u1
Cyp3a11
Cyp3a13
Cyp3a16
Cyp3a25
Cyp3a41a
Cyp3a41b
Cyp3a44
Cyp3a59
Cyp4a10
Cyp4a12a
Cyp4a12b
Cyp4a14
Cyp4f13
Cyp4f14
Cyp4f15
Cyp4v3
Cyp7a1
Cyp7b1
Cyp8b1
Cyp17a1
Cyp26a1
Cyp27a1
Cyp39a1
Cyp51
Counts per million
Layer (central to portal)

*SFig. 7:* ***Cytochrome expression in function of sex, time and space.*** *Cytochromes of the families 1-3, which are the most pharmacokinetically important phase 1 metabolism genes, show high sexual dimorphism. Cyp1a2 shows a strong pericentral bias, and so do Cyp3 family members, including CYP3A4 orthologues. In the case of Cyp2a4 and Cyp2a5, which metabolize xenobiotics, steroids and fatty acids, there is higher expression at ZT10, in anticipation of feeding and detoxification. Both the neutral (classic) and acidic (alternative) pathways of bile acid synthesis are affected, with effects of time and space on Cyp7a1 and high sexual dimorphism in Cyp7b1. Cyp8b1, involved in bile acid hydroxylation, also shows a temporal trend, but also large effects of sex.*

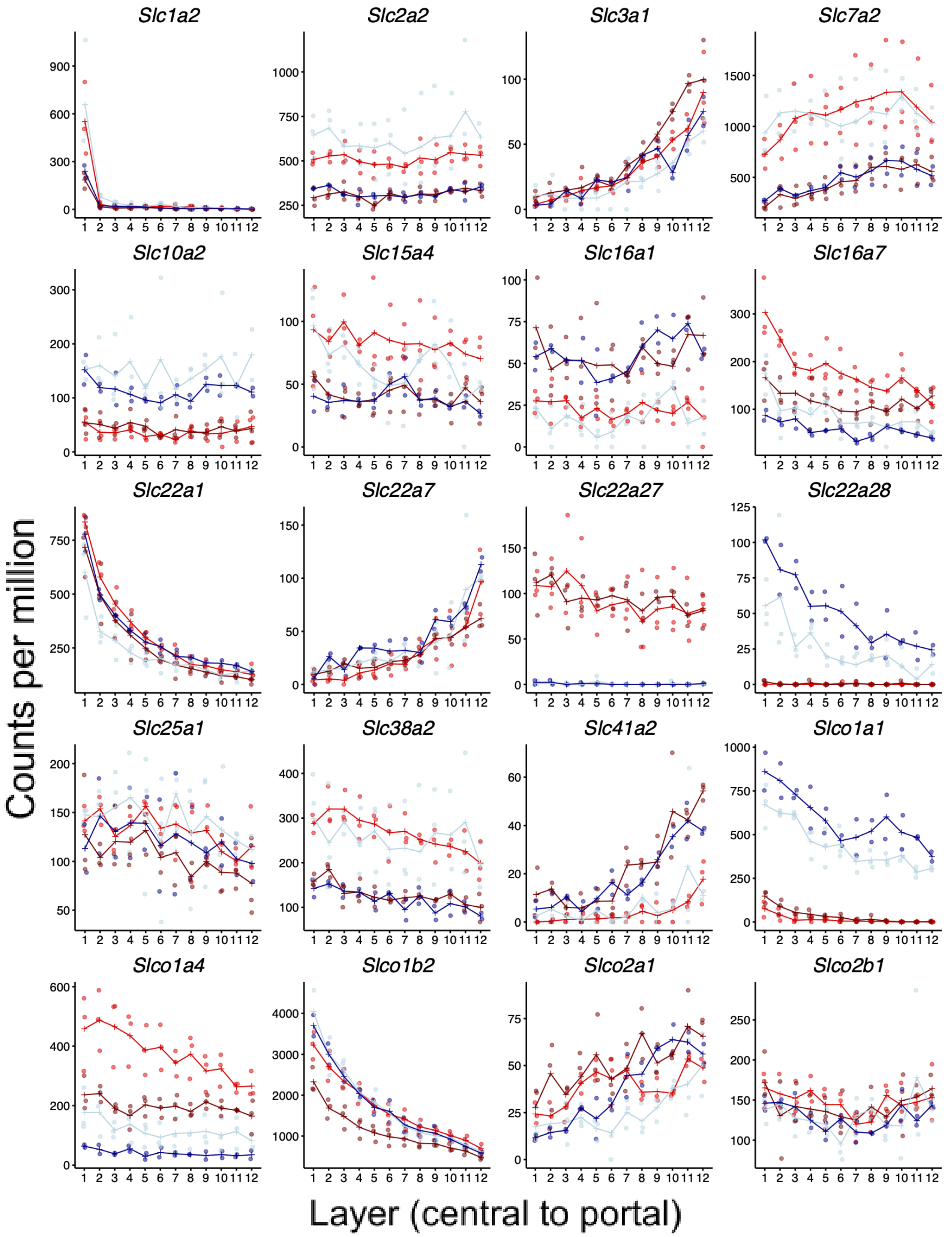


*SFig. 8:* ***Solute carriers.*** *Examples of solute carrier expression. Solute carriers are involved in endo- and xenobiotic biodistribution, including drugs. Slc2a2 encodes GLUT2, is a glucose transporter, with higher expression at the end of the active phase. The amino acid transporter Slc3a1 is much higher periportally. Slc22 family members transport small molecules, metabolites and drugs, and are associated with inherited metabolic disorders and DILI. Slc22a1 encodes OCT1. Slco1a1, which encodes OATP1A1 and is involved in DILI, is much higher in males and shows a pericentral bias. Contrary, Slco1a4, which encodes OATP1A4, is higher in females, and shows higher expression at ZT10.*

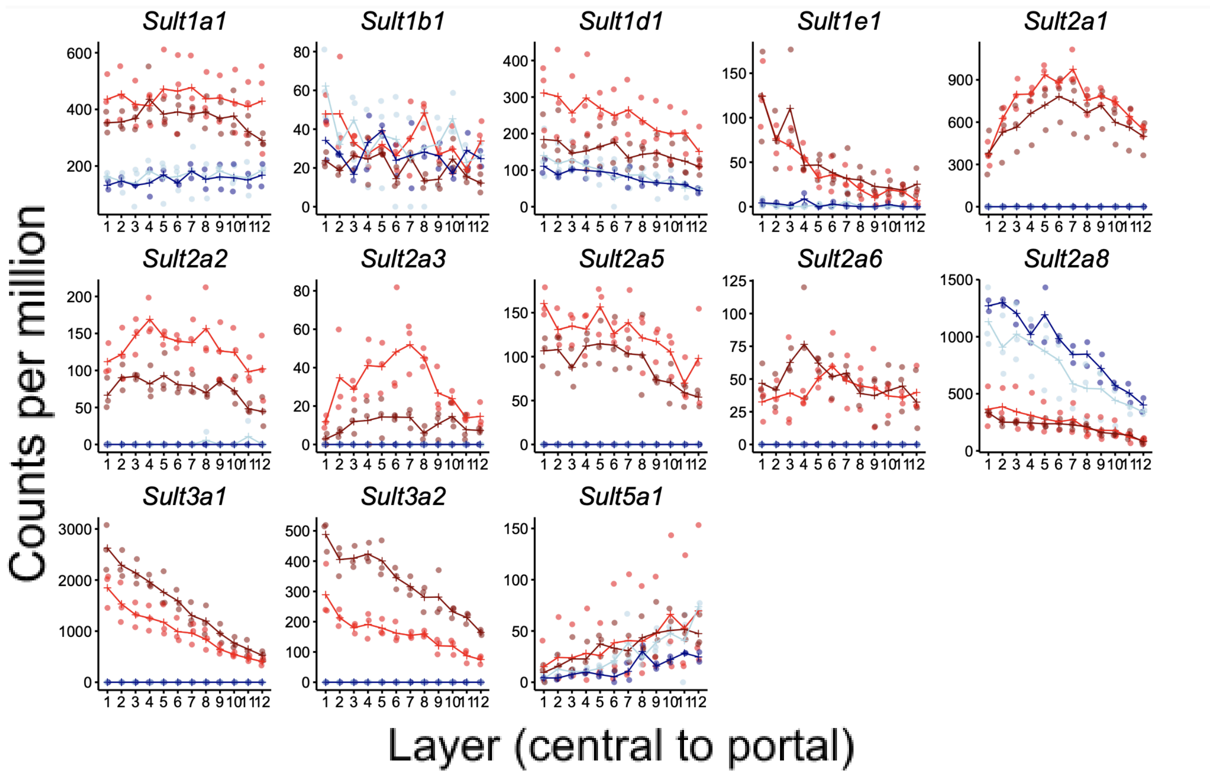


*SFig. 9:* ***Sulfotransferases.*** *Multiple sulfotransferases show a higher expression in females, with the exception of Sul2a8. Sult2a8, higher in males, and Sult3a1, higher in females, are the two sulfotransferases with highest mRNA levels, and both show a pericentral bias.*

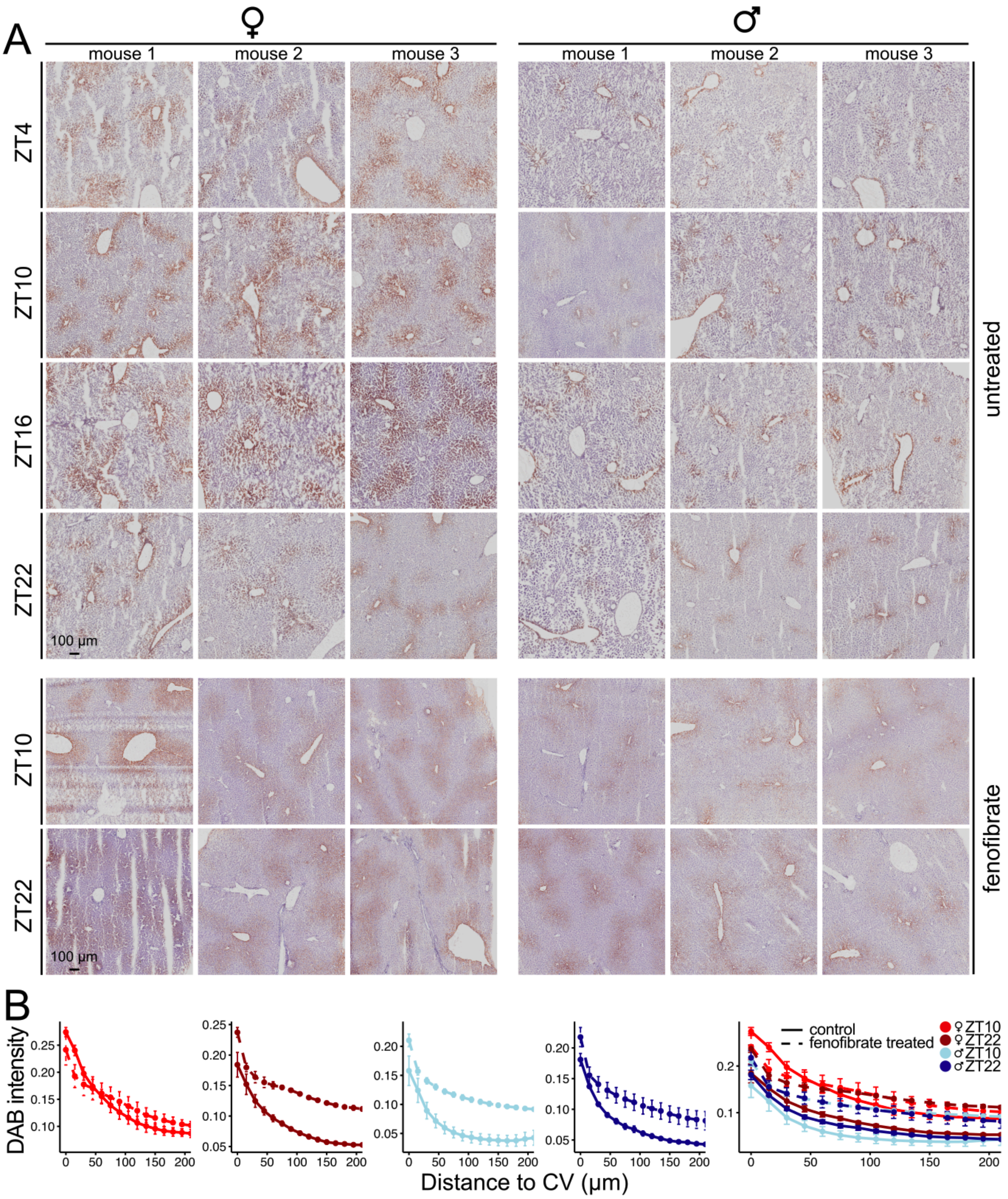


*SFig. 10:* ***H-DAB staining of mouse fresh frozen livers shows a higher immunoreactivity in females and effects of fenofibrate treatment on VLDLR.*** *(A) In all samples, VLDLR immunoreactivity is limited to the pericentral area. In the untreated (control) groups, the immunoreactivity is higher in females, whereas in males the DAB stain appears weaker and less spread. In the treated groups, the spread appears larger compared to the untreated groups, but the VLDLR immunoreactivity in treated males is weaker than the female untreated ZT4 to ZT16 groups. Effects of fenofibrate treatment are sex-dependent. In males, which exhibit only low baseline immunoreactivity, the relative increase in VLDLR upon 5 days of fenofibrate treatment is larger than in females. (B) The quantification of DAB intensity confirms a larger spread and higher immunoreactivity in females, and shows that fenofibrate predominantly increases VLDLR signal further from the central vein, with most profound effects in the male groups and in the female ZT22 group (n = 36; n = 3, per condition; error bars represent the SEM between biological replicates).*

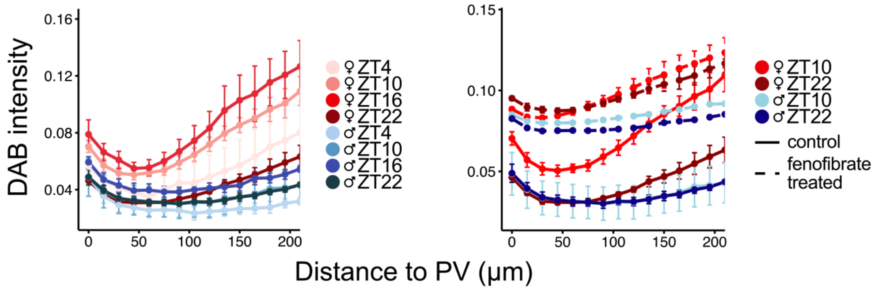


*SFig. 11:* ***Relative H-DAB intensity in function of distance from the portal vein.*** *VLDLR immunoreactivity, based on H-DAB staining around the portal vein (PV). In untreated animals (left panel), the signal increases with distance from the vein, with a higher slope in females at time points ZT10 and ZT16 compared to ZT4 and ZT22, and compared to males. Contrary to immunoreactivity around the central vein, which is high at ZT4, the portal to central ZT4 female slope of DAB intensity is shallow and similar to that at ZT22. Fenofibrate treated animals (right panel) show higher residual immunoreactivity around the portal vein (n = 36; n = 3, per condition; error bars represent the SEM between biological replicates).*

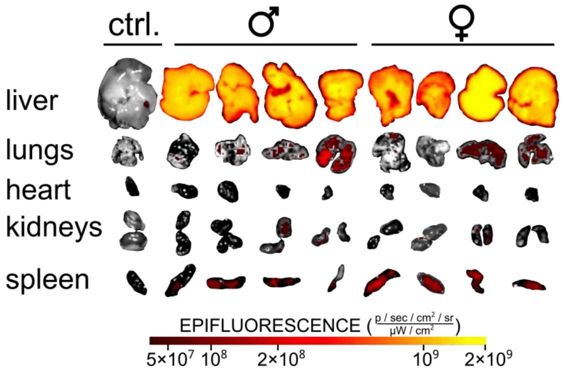


*SFig. 12:* ***Hepatic uptake of fluorescently (DiI) labeled VLDL particles.*** *Purified human fluorescently labeled VLDL particles were injected via the lateral tail vein of living animals. After 20 min, the animals were dissected, and organs flushed of blood. The VLDL-DiI concentrated in the mouse liver at the organ level.*

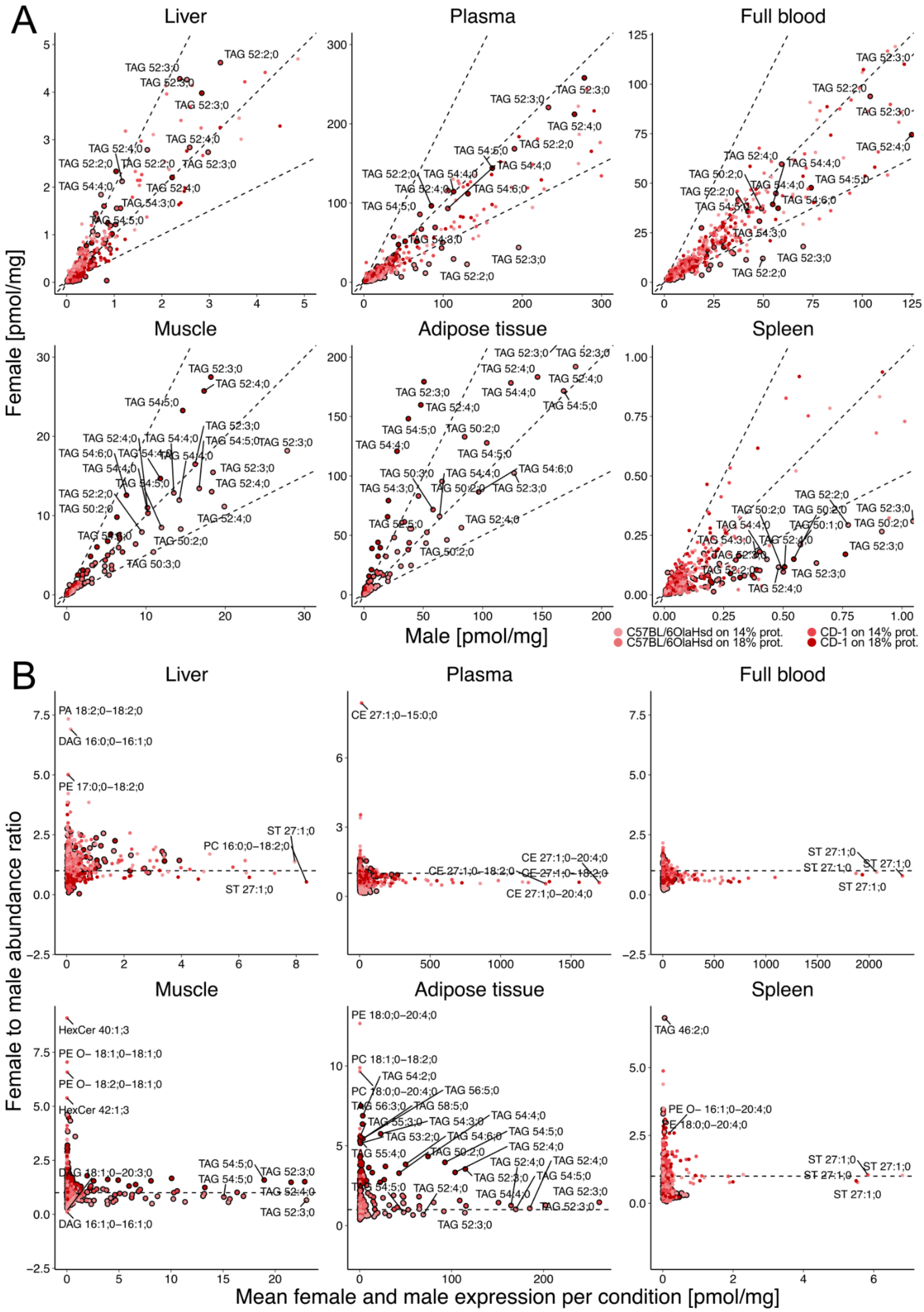


*SFig. 13:* ***Bulk lipidomics in C57BL/6OlaHsd and CD-1 mice (reanalysis of data from ref. 42).*** *The mice were fed a normal chow, 14% protein content diet, or a growth-, gestation- and lactation-supporting 18% protein diet. TAGs are highlighted with black outlines around points. (A) Female and male lipid abundance. Dashed lines*

*show an equivalent (middle line) or two fold difference in abundance (top and bottom lines). Across conditions, the female liver has a higher concentration of the most highly abundant TAGs (TAG 52, TAG 54), whereas in plasma, full blood and the spleen, TAGs are higher in males. (B) Ratio vs. mean abundance plots represent absolute abundance of TAGs compared to other lipids in different tissues.*

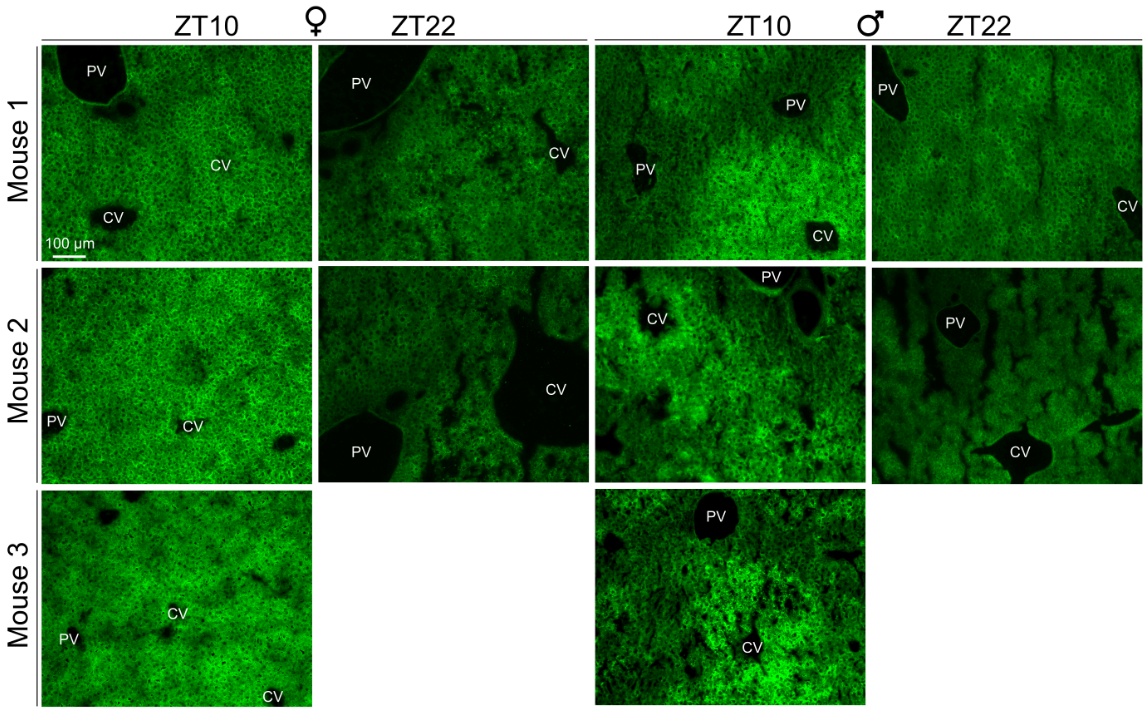


*SFig. 14:* ***BODIPY 493/503 stainings on fresh frozen livers.*** *The neutral lipid dye shows a zonated pattern in male mice at ZT10, but not at ZT22 (n = 2-3, per condition). In females, such intensive zonation is absent.*

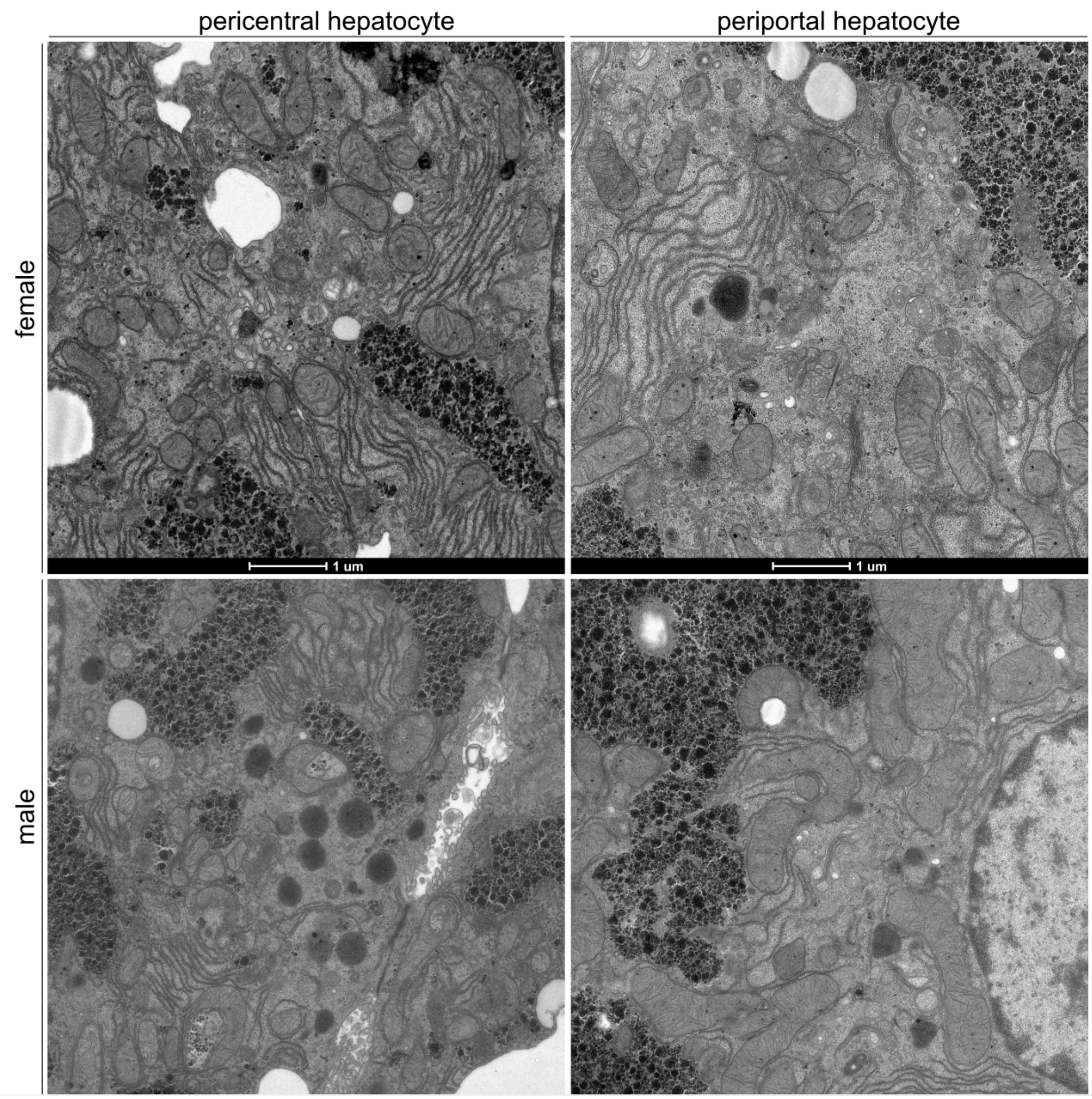


*SFig. 15:* ***Transmission electron microscopy images of mouse liver sections.*** *Vesicles loaded with VLDL particles are abundant periportally. Periportally, mitochondria are larger and more elongated. Biological replicates of mice, shown in the main figure, are shown here.*

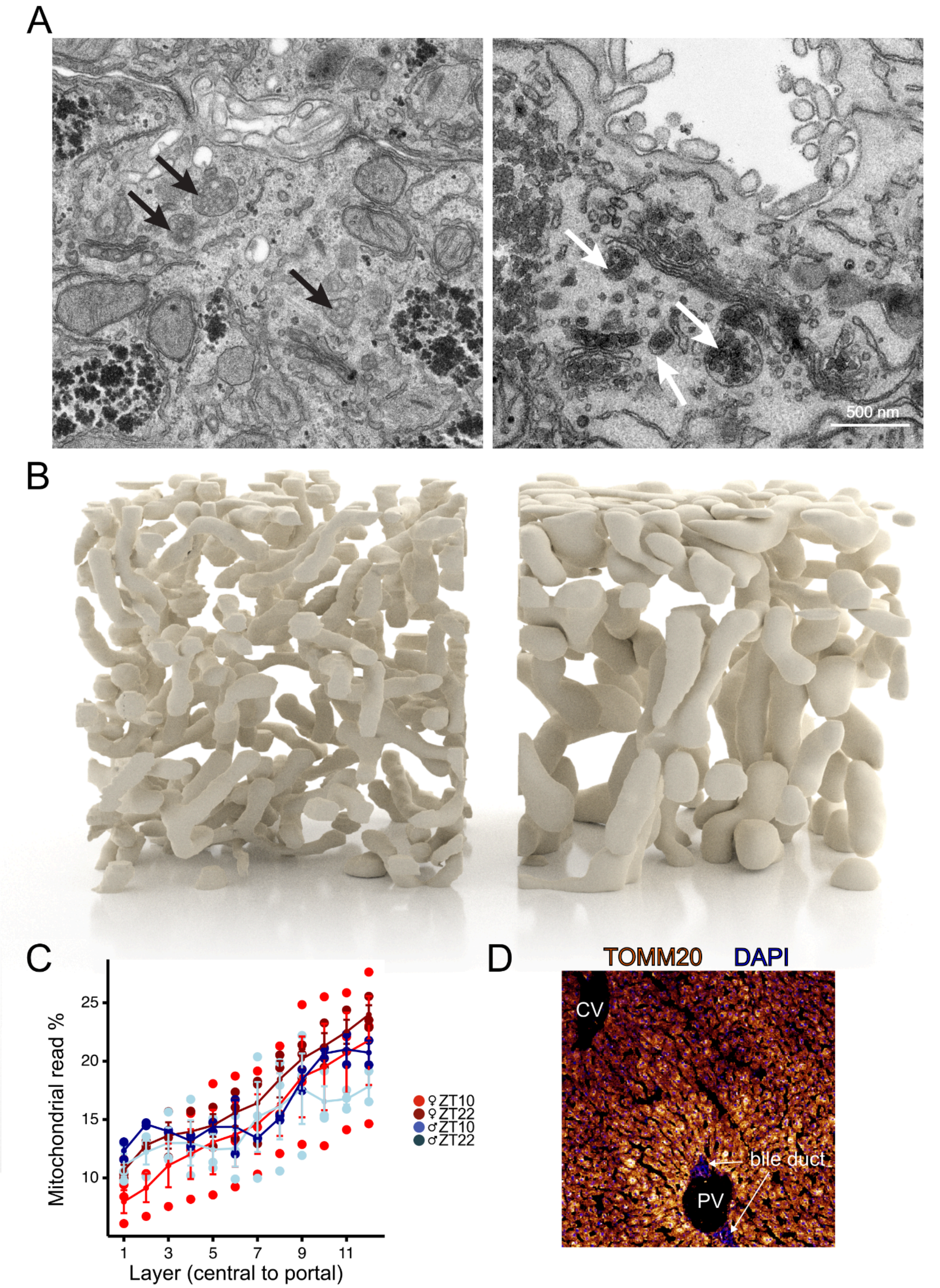


*SFig. 16:* ***Scanning electron microscopy of mouse liver sections and a zone-dependent mitochondrial phenotype.*** *(A) Two different electron microscopy preparation methods render the VLDL particles with different levels of heavy metal stain. The left hand image shows the VLDL carriers (black arrows) stained for transmission*

*electron microscopy (TEM) with a membrane surrounding packets of VLDL particles that show a lightly contrasted center. Whereas the staining method for block face scanning electron microscopy (SEM; right hand image) shows a similar structure with much more darkly stained particles. Scale bar is 500 nm. (B) Mitochondria from two 7 × 7 × 7 μm3 cubes from a male mouse imaged in the central (left), and portal (right) regions. Every mitochondrion was reconstructed in 3D, showing differences in PC vs. PP mitochondrial morphology. (C) The difference in mitochondrial size is reflected in the percent of mitochondrial reads, which increases in the periportal direction (n = 11, mice; dots represent individual cells, error bars are standard errors between biological replicates). (D) A representative TOMM20 (mitochondrial import receptor subunit TOM20 homolog) stain showing higher immunoreactivity in this outer mitochondrial membrane-associated protein in the periportal region. Bile ducts can be identified with DAPI.*

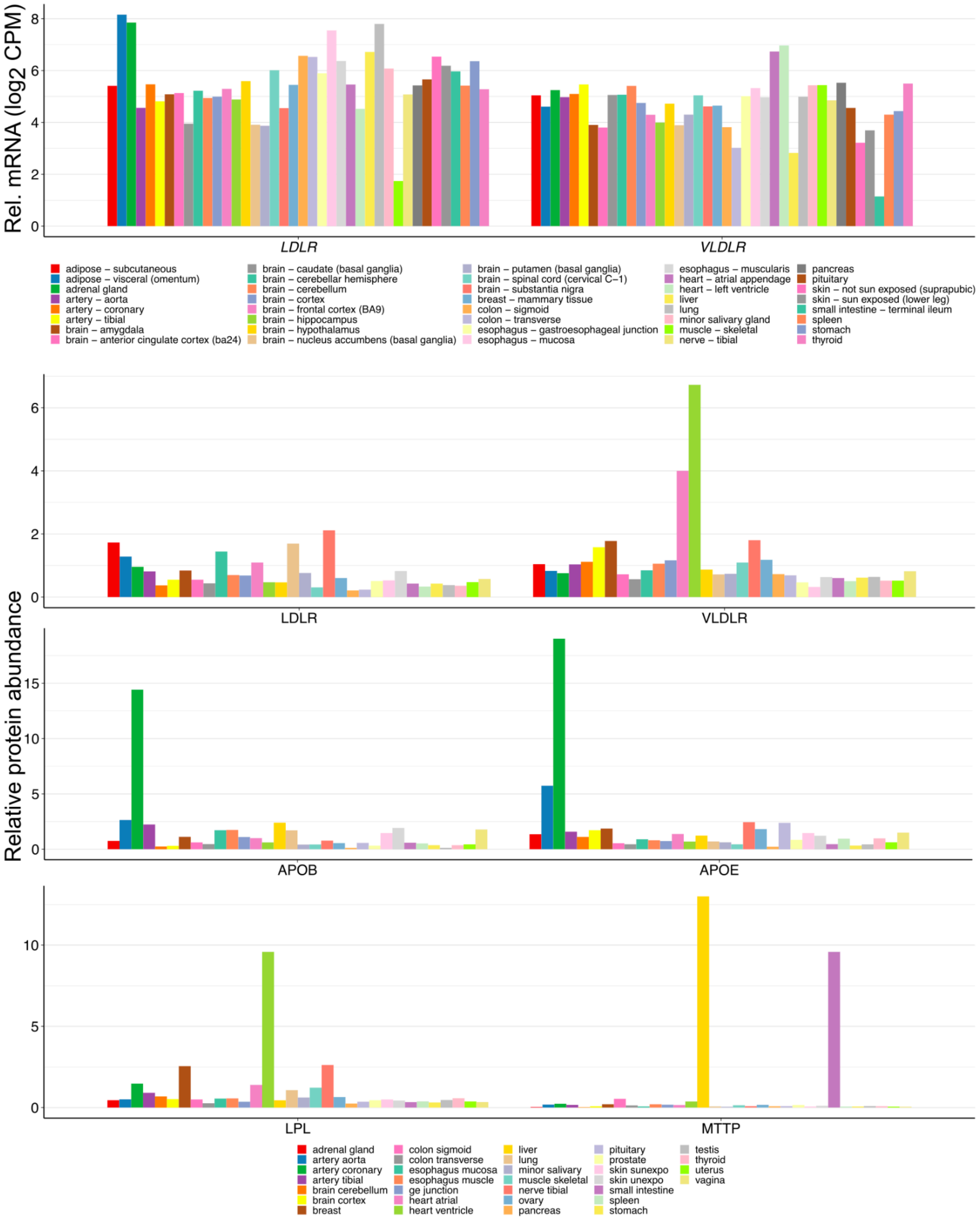


*SFig. 17:* ***Human bulk transcriptome and proteome (reanalysis of proteomics data from ref. 47) across tissues.*** *(A) Bulk mRNA across tissues. Expression of LDLR and VLDLR in bulk human tissue represented as mean values of males and females combined. The lowest VLDLR mRNA levels are in the liver (yellow) and terminal ileum (teal). High LDLR levels are observed in the visceral fat and adrenal gland. (B) Bulk proteome across tissues, median levels are shown. Relative abundance of the VLDLR protein across human tissues, combined values of males and females, suggests that the protein levels do not not correlate to mRNA levels. The highest abundance of the receptor is in the heart. The liver levels are comparable to a multitude of other tissues, despite low RNA levels. Similarly, the measured protein levels of VLDLR are comparable to LDLR in the liver.*

*APOB and APOE levels are highest in the coronary artery, likely pointing to lipoprotein biodistribution. LPL has high specificity, with large abundance in the ventricle, whereas MTTP is liver and small intestine specific.*

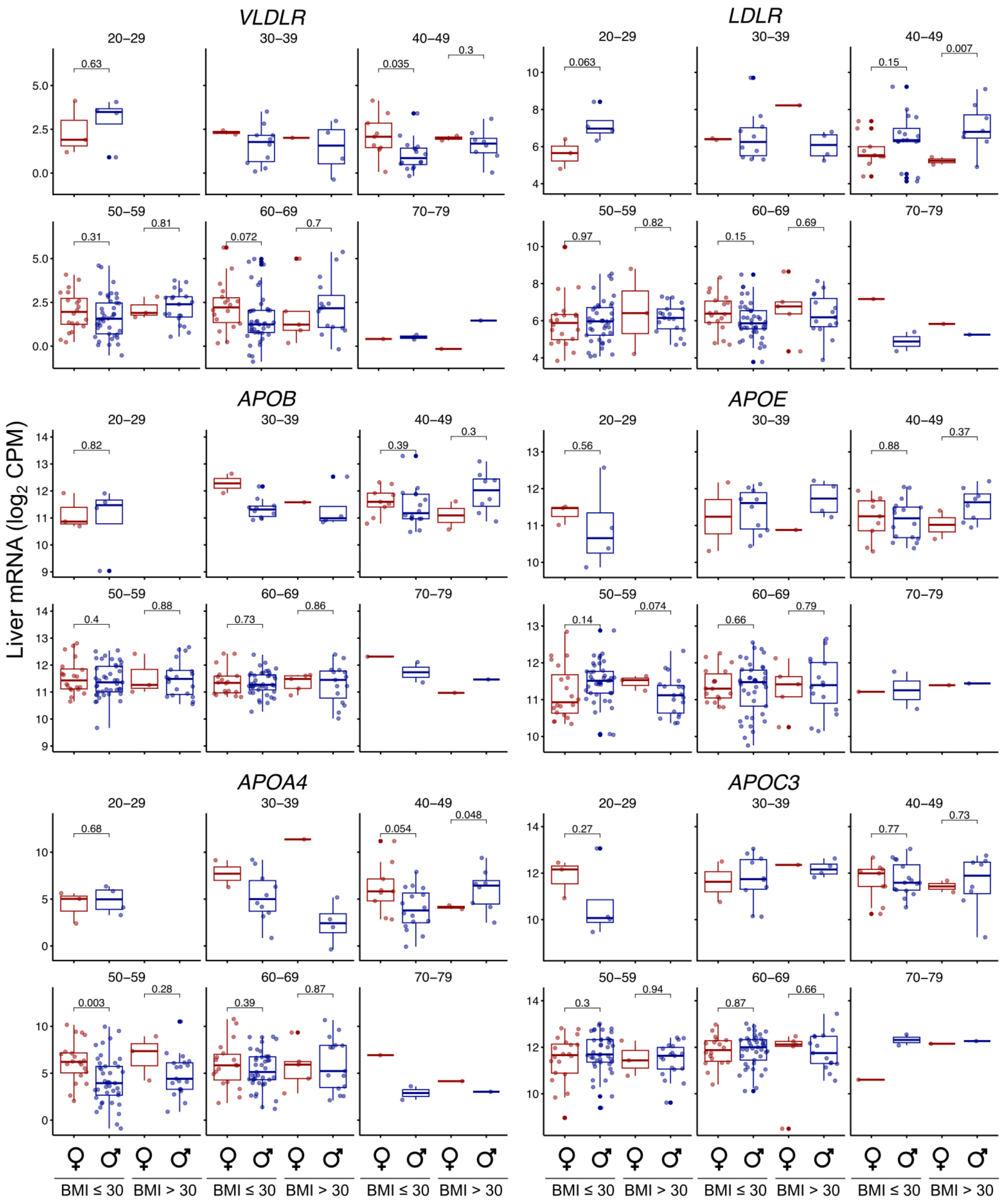


*SFig. 18:* ***Stratification of VLDLR, LDLR and apolipoprotein B, E, A4 and C3 expression in the human liver.*** *The expression was stratified by sex and body mass index (BMI). The center line represents the mean, error bars represent the interquartile range, p-values were calculated with a t-test, without outlier removal (Methods).*

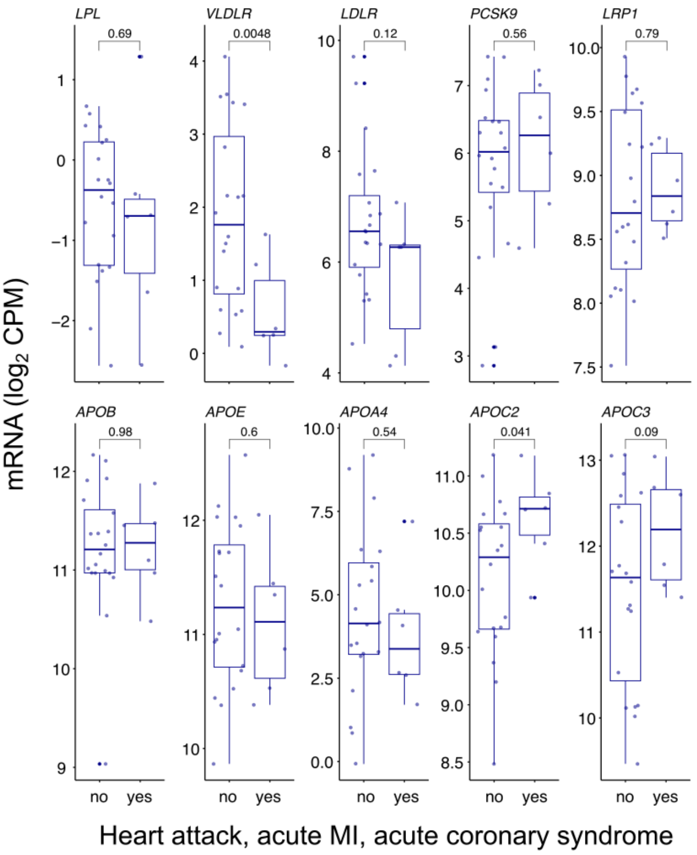


*SFig. 19:* ***Hepatic gene expression in young (≤ 49 years) non-obese (BMI ≤ 30) men.*** *The expression was stratified by whether the donor was annotated with having a heart attack, acute myocardial infarction or acute coronary syndrome (GTEx variable name MHHRTATT in phv00169162.v4.p1). The center line represents the mean, error bars represent the interquartile range, p-values were calculated with a t-test, without outlier removal (Methods). In males that suffered a heart attack, the hepatic VLDLR expression is significantly lower.*

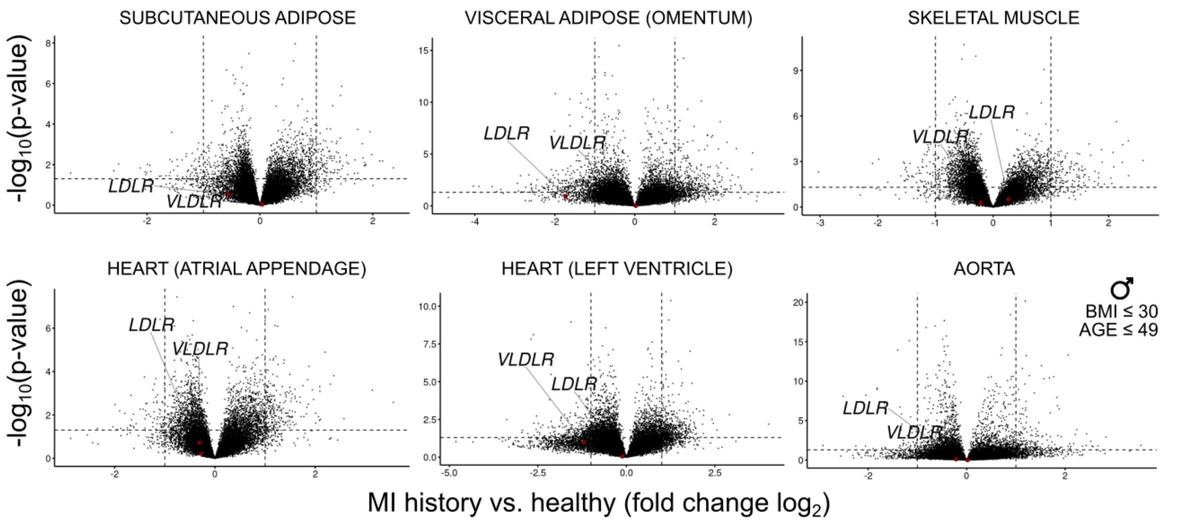


*SFig. 20:* ***Gene expression differences in young non-obese men with or without a medical history of acute myocardial infarction or acute coronary syndrome.*** *Neither VLDLR nor LDLR are significantly differently expressed in donors with such a medical history in the represented non-hepatic tissues. The dashed horizontal line represents the p-value of 0.05.*

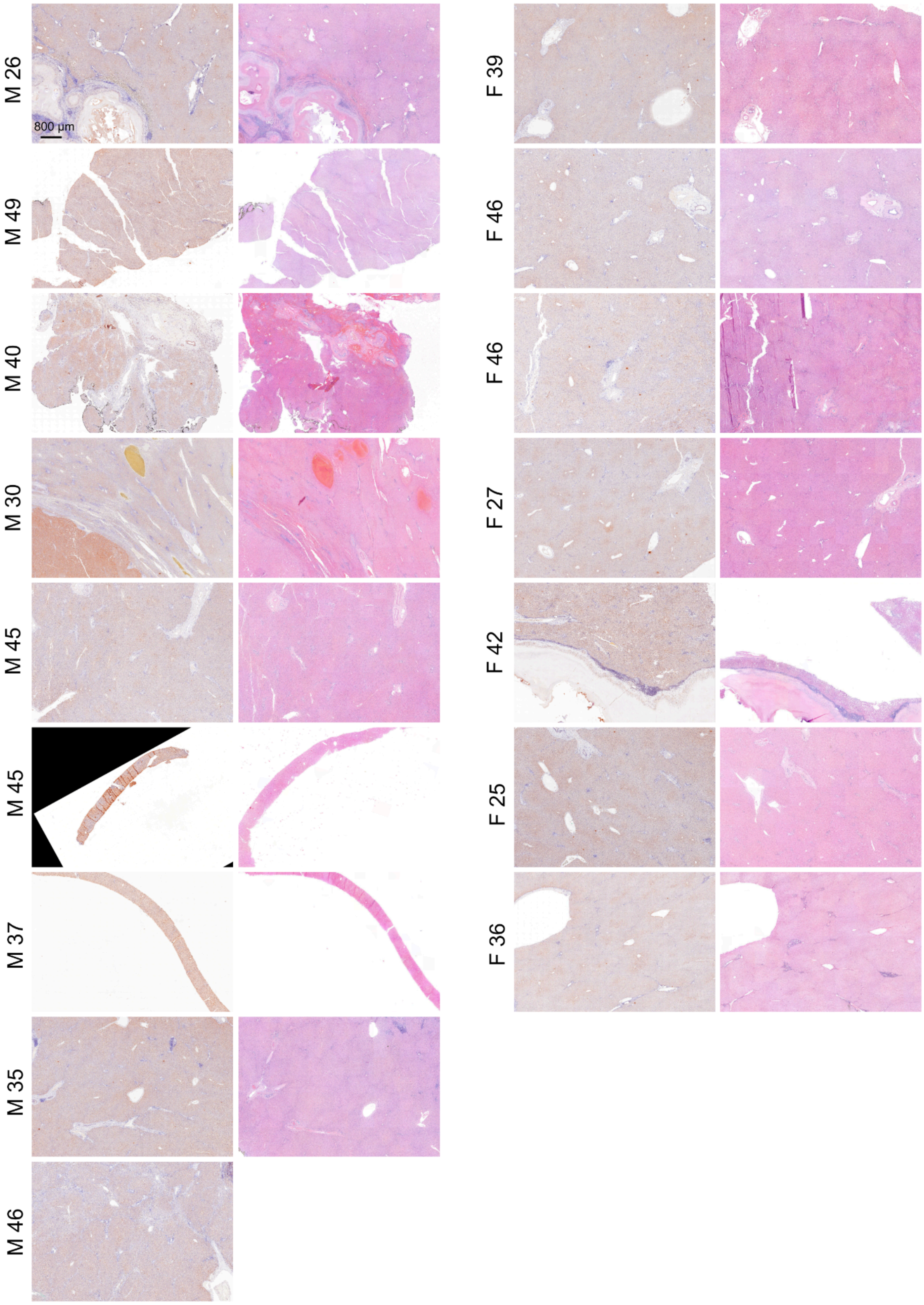


*SFig. 21:* ***Overviews of samples from male and female human donors ≤ 49 years old.*** *Stainings of liver sections of premenopausal women (F) and age-matched men (M) are shown. The hematoxylin and DAB VLDLR staining and the corresponding hematoxylin and eosin staining are shown for each donor, with the age of each*

*donor indicated next to the stained sections. Larger overview areas are shown, representing zonation of VLDLR immunoreactivity and the intraindividual variability. Indications of pathologies are visible, in some cases leading to perturbed zonation.*

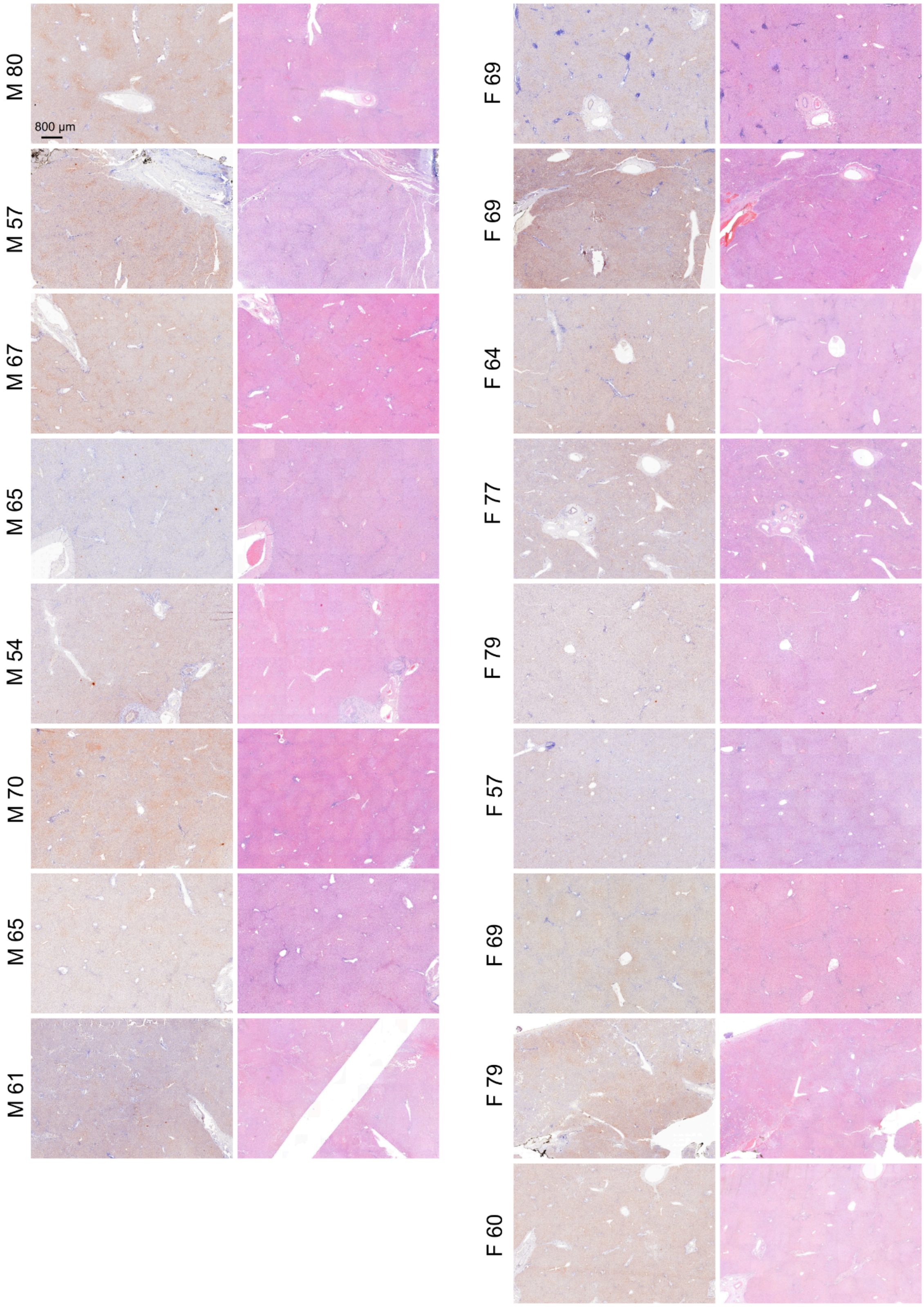


*SFig. 22:* ***Overviews of samples from male and female human donors > 49 years old.*** *Stainings of liver sections of postmenopausal women (F) and age-matched men (M) are shown. The hematoxylin and DAB VLDLR staining and the corresponding hematoxylin and eosin staining are shown for each donor, with the age of each*

*donor indicated next to the stained sections. Larger overview areas are shown, representing zonation of VLDLR immunoreactivity and the intraindividual variability. Indications of pathologies are visible, in some cases leading to perturbed zonation.*

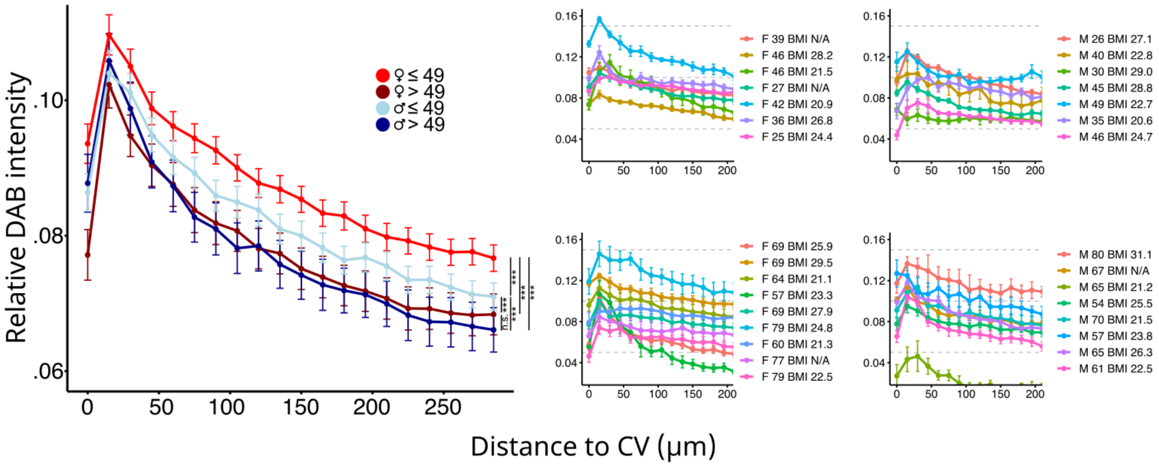


*SFig. 23:* ***Quantified relative intensity of DAB staining in human sections.*** *(A) Mean relative DAB signal per group, donors were grouped by age and sex as shown in the legend (n = 31; errors represent the SEM between individual samples within a group). The staining in the groups of men and women > 49 years are potentially overestimated due to the presence of the age pigment lipofuscin. (B) Relative intensity of DAB staining on a per sample basis, errors represent the SEM between veins.*

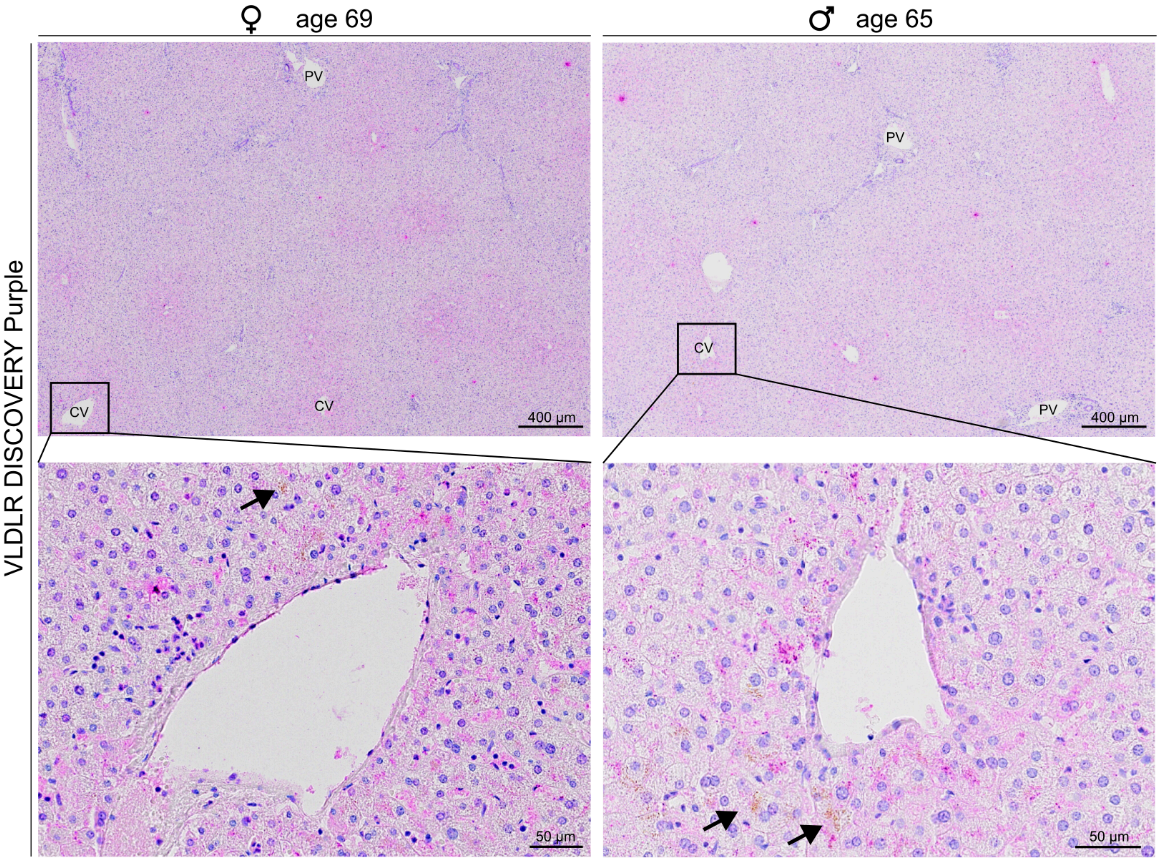


*SFig. 24:* ***VLDLR stainings in purple and fluorescence.*** *Representative hematoxylin and VLDLR stainings with immunoreactivity revelation in purple. In both the female donor (left) as well as the male donor (right), stronger immunoreactivity is observed around pericentral veins (upper panels). In the higher magnification view (bottom panels), the arrows point to lipofuscin. Despite a strong presence of this age pigment in these representative samples of older donors, the purple immunoreactivity is dominant, supporting the fact that a large proportion of the signal in H-DAB stainings comes from immunoreactivity. The DISCOVERY Purple chromogen was used. CV - central vein, PV - portal vein.*

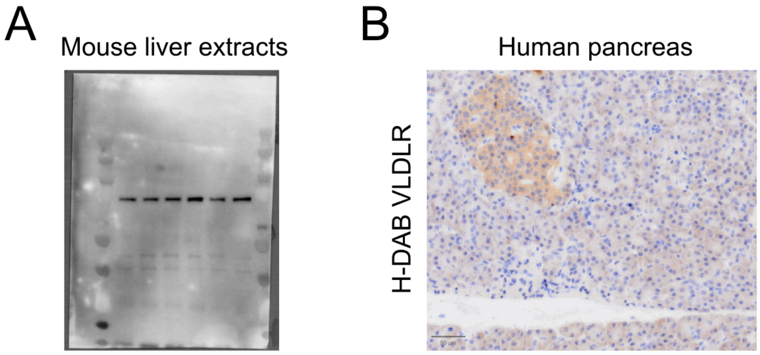


*SFig. 25:* ***VLDLR antibody validation.*** *The antibody was tested on (A) mouse liver extracts in a western blot, and on (B) human FFPE pancreatic sections. The antibody consistently produces a single band in the western blot and in the pancreas it stains the islets. These immunostainings were not optimized.*
